# Cell therapy for regeneration of injured donor lungs for transplantation

**DOI:** 10.64898/2026.03.16.712049

**Authors:** Franziska Olm, Margareta Mittendorfer, Dag Edström, Anna Niroomand, Nicholas B. Bèchet, Gabriel Hirdman, Haider Ghaidan, Embla Bodén Janson, Michaela Oeller, Katharina Schallmoser, Gunilla Kjellberg, Martin Stenlo, Stefan Scheding, Snejana Hyllén, Sandra Lindstedt

## Abstract

Donor organ shortage remains the major barrier to transplantation resulting in deaths on the waiting list. For lungs, aspiration-related injury is a common cause of donor organ discard and increases the risk of primary graft dysfunction. Currently, no effective therapies exist to repair damaged donor lungs prior to transplantation. Here, we investigated whether mesenchymal stromal cells (MSCs) from bone marrow or full-term amniotic fluid could restore severely injured donor lungs in a porcine model integrating ex vivo lung perfusion, transplantation and post-transplant follow-up (n=48; 24 donors, 24 recipients). MSCs were administered either once during ex vivo lung perfusion or repeatedly across lung perfusion and the early post-transplant period and compared with placebo treated controls. A single dose conferred only partial benefit, whereas repeated dosing restored graft function, normalized gas exchange and haemodynamics, and prevented graft dysfunction. MSCs from both sources were similarly effective in repeated regimens. These findings identify dosing schedule, rather than cell source, as key determinant of durable organ rescue and support perfusion-guided cell therapy as potentially generalizable regenerative strategy across solid-organ transplantation.

## Introduction

Lung transplantation (LTx) is the only curative treatment for end-stage pulmonary disease^1^, but its impact is limited by donor organ shortage and underuse of available grafts. Up to 80% of donor lungs are deemed unsuitable for transplantation, often because of acute lung injury caused by aspiration, infection, or neurogenic pulmonary oedema, which can progress to acute respiratory distress syndrome (ARDS)^2,3^. The World Health Organization has classified the global shortage of transplantable organs as a public health crisis, with transplantation activity fulfilling only approximately 10% of the demand^4^. Currently, no effective treatment strategies exist to recover severely injured donor lungs and make them suitable for transplantation. Such grafts are closely linked to primary graft dysfunction (PGD), the leading cause of early post-transplant mortality. PGD affects approximately one-third of recipients and substantially increases the risk of chronic lung allograft dysfunction (CLAD), the major cause of late graft failure and long-term mortality^2,5–8^.

Ex vivo lung perfusion (EVLP), originally developed at Lund University^9,10^ and subsequently advanced into clinical practice by the Toronto Lung Transplant Program^11,12^, is increasingly used to reassess marginal or previously discarded donor lungs prior to implantation. By maintaining donor lungs under normothermic ventilation and perfusion in a controlled extracorporeal setting, EVLP enables real-time functional assessment and has helped expand the donor pool clinically^11^. Beyond assessment, EVLP provides a therapeutic window for direct delivery of regenerative interventions to the isolated organ^13^. Although such applications remain largely preclinical, studies have demonstrated the potential of EVLP for organ repair and immune modulation^14,15^, notably through the administration of mesenchymal stromal cells (MSCs)^16–20^. MSCs exhibit broad immunomodulatory and tissue-protective properties, interacting with both innate and adaptive immune systems^21–23^, and their low immunogenicity supports their use in transplantation^19^. However, important translational questions remain unresolved. MSCs can be derived from multiple tissue sources, and although preclinical studies suggest functional differences among them^24–26^, comparative functional *in vivo* data are scarce, especially in LTx, and generally lack post-transplantation evaluation^27^. Early-phase clinical studies of bone marrow- and adipose tissue-derived MSCs in lung transplantation have shown feasibility, but efficacy signals remain inconclusive^28–31^. Moreover, MSC therapy has not been evaluated for restoring aspiration-damaged donor lungs in a transplant setting, as preclinical studies have mainly focused on ischemia-reperfusion injury (IRI) during EVLP^16,17,32,33^.

Building on our previous findings that repeated dosing of full-term amniotic fluid-derived MSCs (TAF-MSCs) can mitigate infection in donor lungs and improve recipient outcomes^20^, we evaluated MSC therapy to restore aspiration-damaged donor lungs in a porcine transplantation model. TAF-MSCs and bone marrow-derived MSCs (BM-MSCs) were compared, and both single-dose administration during EVLP and repeated dosing across EVLP and post-transplantation were assessed. Recipients were allocated to four groups: non-treated controls, single-dose BM-MSCs, repeated-dose BM-MSCs, or repeated-dose TAF-MSCs. We hypothesized that repeated MSC dosing would provide greater and more durable benefit than a single intra-EVLP dose, and that comparison of two MSC sources would clarify whether tissue origin or dosing schedule is the principal determinant of efficacy. Our findings identify dosing schedule, rather than cell source, as a key determinant of durable regenerative efficacy and support cell therapy as a strategy to rescue severely injured donor lungs and improve outcomes.

## Results

### Establishment of aspiration-induced lung injury

Administration of acidified gastric contents (pH 2) reproducibly induced severe donor lung injury while preserving overall haemodynamic stability with minimal inotropic support (Fig. 1 and Table 1). Injury was marked by increased pulmonary vascular resistance and systemic vascular resistance, reduced dynamic compliance, and a fall in PaO_2_/FiO_2_ from 521.0±44.3 at baseline to 193.7±57.2 after injury induction (all p<0.0001; Fig. 2a). All donors developed radiographic infiltrates, bronchoscopic signs of airway inflammation, and gross features of acute lung injury (Fig. 2b and Extended Data Fig. 1a–c).

**Fig. 1:**
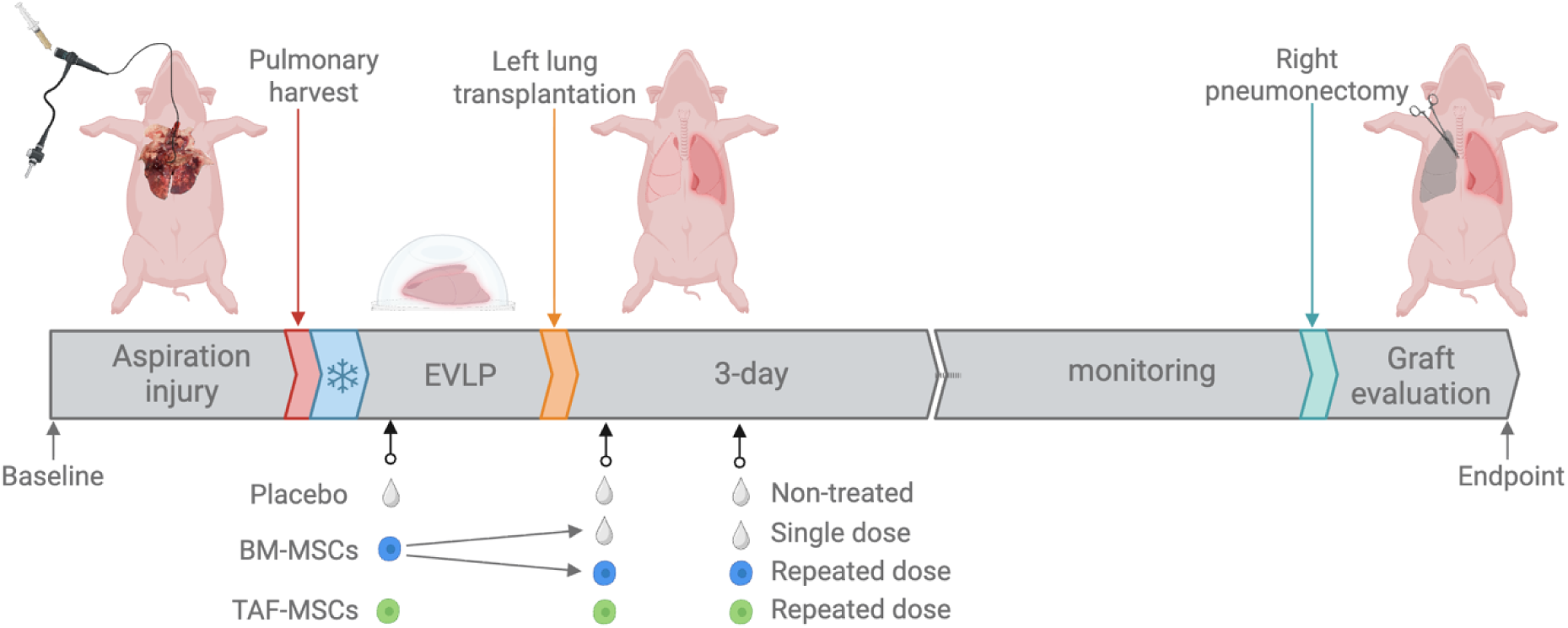
Experimental design of MSC therapies in lung transplantation. Timeline of aspiration-induced lung injury and administration of mesenchymal stromal/stem cells (MSCs) or placebo during *ex vivo* lung perfusion (EVLP) and post lung transplantation (LTx). Lung injury was established over 6 hours in donor pigs (adult, Yorkshire, n=24) using gastric content with pH 2. Injured lungs were harvested, subjected to 2 hours of cold ischemic time, and 4 hours of EVLP. A left lung transplantation was then performed, and recipients (n=24) were maintained under anaesthesia and monitored for 72 hours. After right pneumonectomy, the function of the transplanted left lung was assessed for an additional 4 hours in isolation. Experimental pairs were randomized to non-treated (placebo, n = 6, white drops) or MSC therapy groups: single-dose bone marrow–derived MSCs (BM-MSCs, n = 6; blue), repeated BM-MSCs (n = 6), or repeated term amniotic fluid–derived MSCs (TAF-MSCs, n = 6; green). MSCs or placebo were delivered during EVLP and post-LTx. Throughout the experiment, haemodynamic and ventilatory parameters, arterial blood gases, plasma/EVLP perfusate, bronchoalveolar lavage, and tissue biopsies were collected for downstream analysis. Figure created with Biorender.com.

**Fig. 2:**
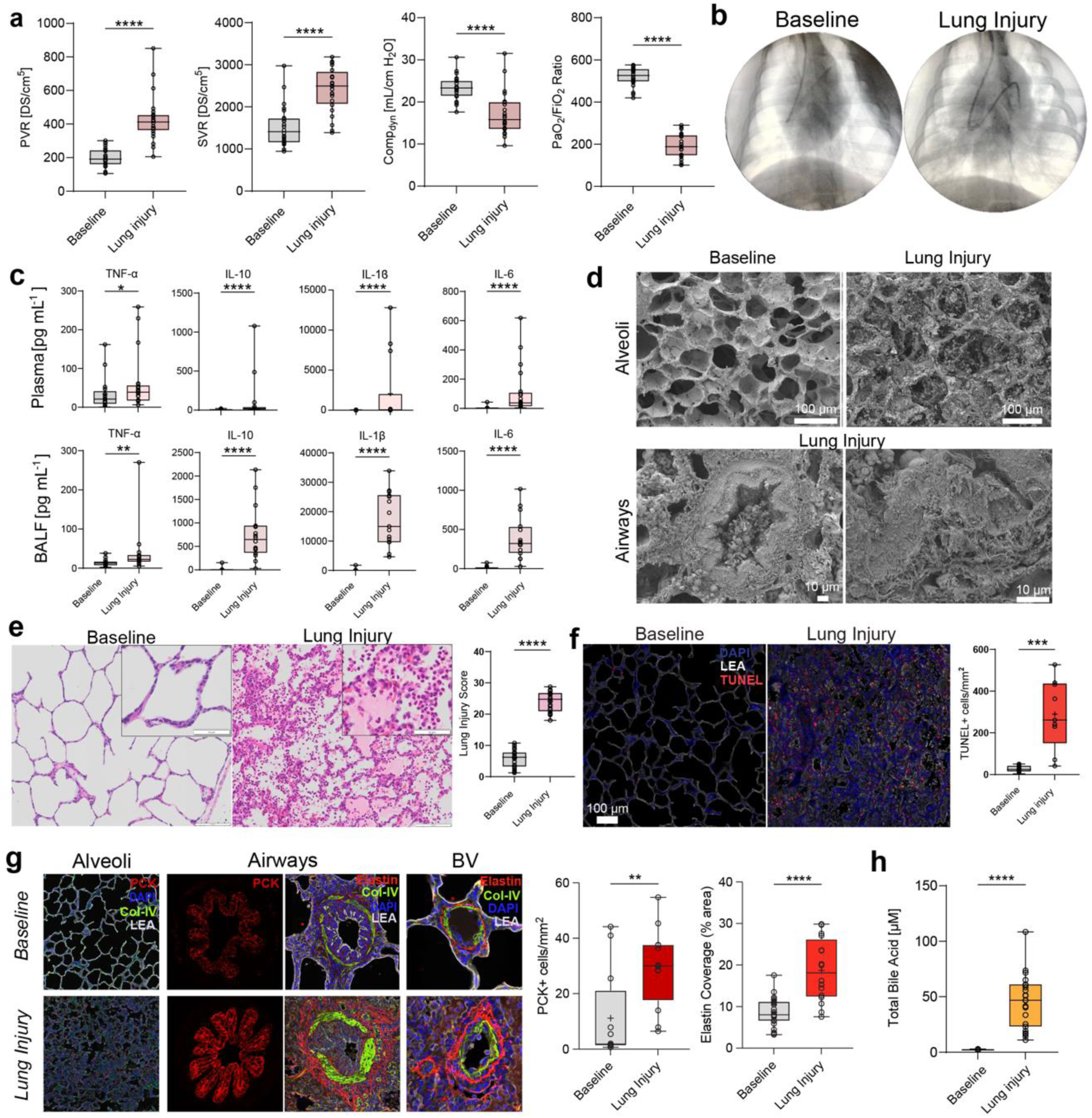
Induction and confirmation of aspiration-induced lung injury in the donors. Gastric content (pH 2) was instilled into donor lungs (n=24), and pigs were observed for six hours under anaesthesia. **a)** Hemodynamic and respiratory parameters at baseline (light grey) and after lung injury (dark pink), including pulmonary vascular resistance (PVR), systemic vascular resistance (SVR), dynamic compliance (Comp_dyn_), and PaO₂/FiO₂ ratio. **b)** Representative chest x-rays at baseline and confirmed lung injury. **c)** Cytokine levels at baseline and after lung injury in plasma (n=17 and n=22, respectively, upper) and bronchoalveolar lavage fluid (BALF, n=16 per group, lower). **d)** Scanning electron microscopy (SEM) images of alveoli (baseline vs lung injury) and airways at lung injury. Scale bars indicate 100 µm (upper) and 10 µm (lower). **e)** Representative haematoxylin and eosin (H&E) stained lung sections at baseline (left) and after injury (middle), scale bars represent 100 µm (overview) and 50 µm (callout). Quantification of histological lung injury scoring (right). **f)** Terminal deoxynucleotidyl transferase dUTP nick end labelling (TUNEL, red), 4′,6-diamidino-2-phenylindole (DAPI, blue), and Lycopersicon esculentum lectin (LEA), white at baseline (left) and at confirmed lung injury (middle). Scale bar represents 100 µm. Quantification of TUNEL^+^ cells/mm^2^ (n=9, right). **g)** Immunostaining for pan-cytokeratin (PCK, red) in alveoli (left) and airways (second to left), elastin in airways (second to right), and in blood vessels in baseline (upper) and at lung injury (lower). 4′,6-diamidino-2-phenylindole (DAPI, blue), collagen IV (Col-IV, green), and Lycopersicon esculentum lectin (LEA, white) staining for visualization of tissue scaffold. Quantification of PCK^+^ cells/mm^2^ (n=12, left) and elastin coverage of vessels (% area, n=23 baseline, n=16 lung injury, right). **h)** Total bile acids (TBA) in BALF at baseline (n=6) and at confirmed lung injury (n=24). Statistical differences were assessed using Student t-test or the Mann-Whitney U-test, where data were not normally distributed. *p<0.05, **p<0.01, ***p<0.001, ****p<0.0001. Boxplots: centre line, median; box, interquartile range; whiskers, min-max. If not otherwise stated, all animals were included in statistical analyses.

**Table 1:** Clinically relevant measurements at baseline and after establishment of aspiration-induced lung injury in donors.

|  | Baseline | Lung Injury |
| --- | --- | --- |
| Sat (%) | 98.8±1.2 | 94.9±3.2 |
| HR (bpm) | 89.8±21 | 89.2±12.3 |
| SBP (mmHg) | 114±15.6 | 115.2±15.4 |
| DBP (mmHg) | 72.8±13.2 | 85.8±12.9 |
| MAP (mmHg) | 87±12.5 | 98.7±13 |
| CVP (mmHg) | 5.8±3.3 | 6.6±2.8 |
| SPAP (mmHg) | 24.3±5.6 | 33.6±6.8 |
| DPAP (mmHg) | 12.8±4.5 | 20.2±5.6 |
| MPP (mmHg) | 17.8±4.5 | 25.8±6.2 |
| PAPWP (mmHg) | 7.8±4 | 10.4±3.2 |
| CO (L/min) | 4.2±0.9 | 3.2±0.7 |
| CI (L/min/m<sup>2</sup>) | 3.8±0.8 | 2.7±0.6 |
| SVRI (Dynes*s/cm<sup>5</sup>*m<sup>2</sup>) | 1833.6±582 | 2909±657.3 |
| PVRI (Dynes*s/cm<sup>5</sup>*m<sup>2</sup>) | 233.3±66.6 | 486.6±123.8 |
| pH | 7.4±0 | 7.4±0 |
| PaCO<sub>2</sub> (kPa) | 6±0.6 | 6.7±0.6 |
| Hb (g/L) | 103.8±9.8 | 110.2±12.6 |
| Lactate (mmol/L) | 1.1±0.2 | 1.5±0.7 |
| BE (mmol/L) | 5.6±2 | 4.3±2.5 |
| MV (L/min) | 6.3±0.9 | 7±1.2 |
| PIP (cmH<sub>2</sub>O) | 17.3±1.5 | 22.5±3.9 |
| PEEP (cm H<sub>2</sub>O) | 5.1±0.3 | 5±0 |
| Vt (mL) | 282.3±23 | 281.7±28.7 |
| RR (breaths/min) | 21.7±2.3 | 24.7±3.6 |

All parameters are presented as mean ± SD values: oxygen saturation (Sat, %), heart rate (HR, beats per minute, bpm), systolic blood pressure (SBP, mmHg), diastolic blood pressure (DBP, mmHg), mean arterial pressure (MAP, mmHg), central venous pressure (CVP, mmHg), systolic pulmonary pressure (SPAP, mmHg), diastolic pulmonary pressure (DPAP, mmHg), mean pulmonary pressure (MPP, mmHg), pulmonary artery wedge pressure (PAPWP, mmHg), cardiac output (CO, L/min), cardiac index (CI, L/min/m<sup>2</sup>), systemic vascular resistance index (SVRI, Dynes\*s/cm<sup>5</sup>\*m<sup>2</sup>), pulmonary vascular resistance index (PVRI, Dynes\*s/cm<sup>5</sup>\*m<sup>2</sup>), pH, partial pressure of carbon dioxides (PaCO<sub>2</sub>, mmHg), haemoglobin (Hb, g/L), lactate (mmol/L), base excess (BE, mmol/L); mechanical ventilator settings with volume-controlled ventilation: minute volume (MV, L/min), peak inspiratory pressure (PIP, cmH<sub>2</sub>O), peak end expiratory pressure (PEEP, cmH<sub>2</sub>O), tidal volume (Vt, mL), respiratory rate (RR, breaths/min).

**Table 2:**
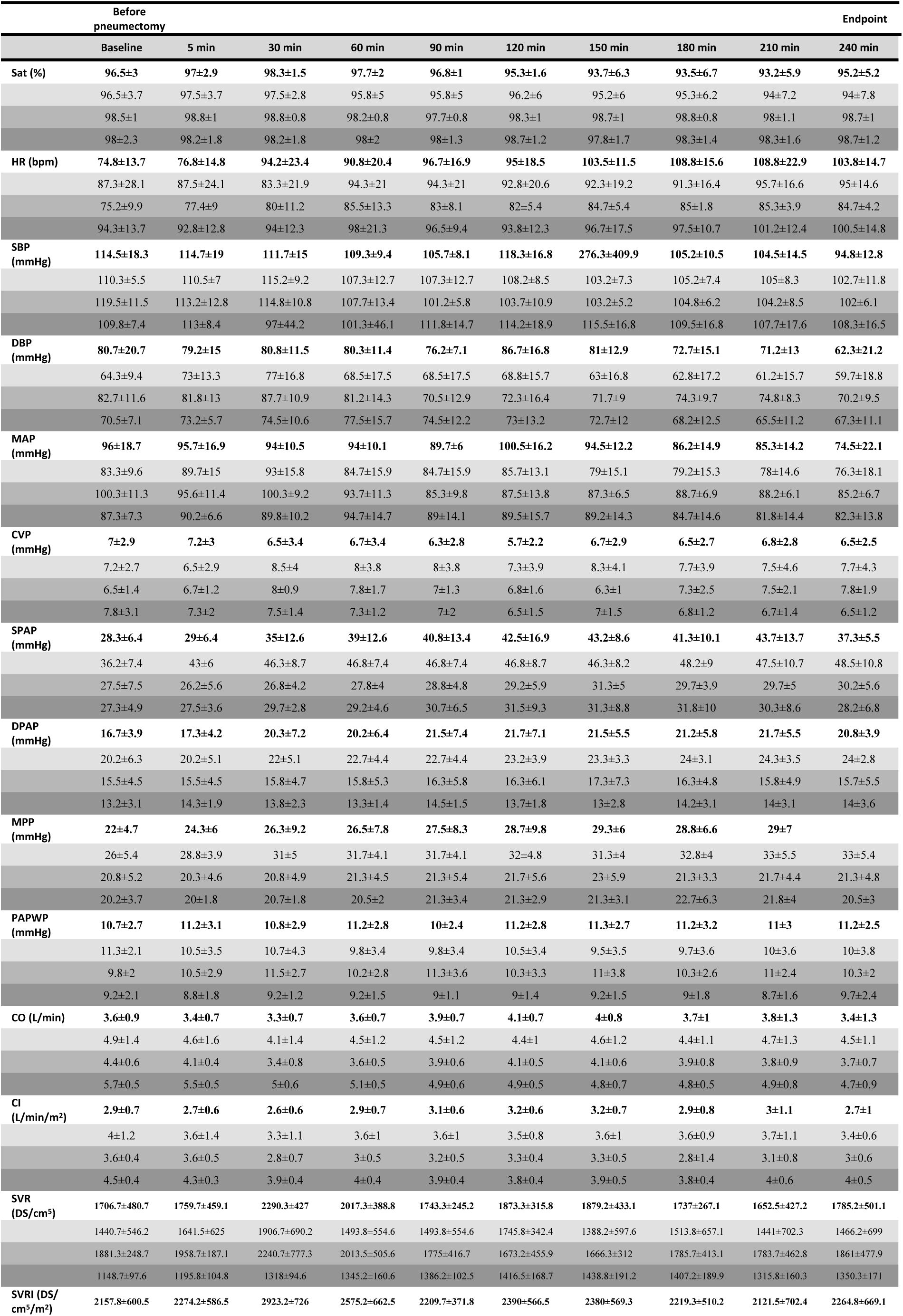
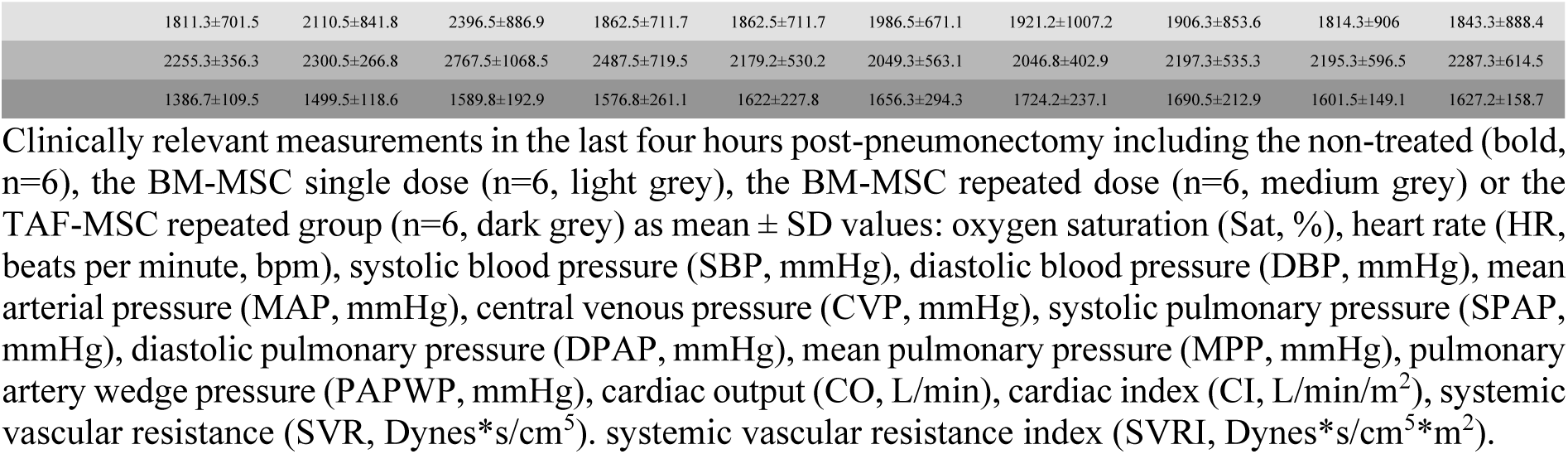
Clinically relevant parameters before pneumectomy and during the four last hours.

Aspiration injury was accompanied by a pronounced inflammatory response. Cytokine analysis revealed significant increases in both plasma and bronchoalveolar lavage fluid (BALF), including TNF-α, IL-10, IL-1β, IL-6, IL-8, IFN-α, IFN-γ, IL-4 and IL-12 (Fig. 2c and Extended Data Fig. 1d), consistent with robust local and systemic immune activation.

Structural analyses confirmed extensive tissue injury. Volumetric scanning electron microscopy and high-resolution confocal imaging showed preserved alveolar architecture at baseline, whereas injured lungs exhibited disrupted alveolar walls, immune cell infiltration, loss of bronchial cilia, airway wall injury, and large cell–neutrophil extracellular trap clusters (Fig. 2d and Extended Data Fig. 1e). Histopathological scoring confirmed significantly increased lung injury after aspiration, demonstrating immune cell infiltration and pulmonary oedema (p<0.0001; Fig. 2e and Extended Data Fig. 1f). Terminal deoxynucleotidyl transferase-mediated dUTP nick-end labelling (TUNEL) staining demonstrated increased apoptosis after injury induction (p=0.0002, Fig. 2f). Airway epithelial disruption and airway and vascular structural changes were assessed by pan-cytokeratin (PCK), smooth-muscle actin (SMA) and elastin staining (Fig. 2g). Compared with baseline, airway PCK expression and vascular elastin coverage were both profoundly upregulated after injury (% area, p<0.0001). The inflammatory response was further supported by a 50-fold increase in total bile acids in BALF (p<0.0001; Fig. 2h), an increase of 13.7×10⁶ total cells/mL in BALF (p=0.0001, Extended Data Fig. 1g), and increased white blood cell (p=0.0303) and neutrophil counts (p=0.0007) in whole blood (Extended Data Fig. 1h). Together, these findings establish a robust aspiration-induced lung injury model meeting Berlin criteria for ARDS, after which lungs were harvested en bloc and cold-stored for two hours^34^.

### Improved lung function, inflammation and morphology following MSC treatment during EVLP

After two hours of cold ischaemic storage in Perfadex^®^ PLUS solution, mimicking clinical transplantation, all grafts were connected to a cellular EVLP system for four hours and randomized to one of three treatment groups: BM-MSC treatment (n=12), TAF-MSC treatment (n=6), or placebo (n=6, non-treated). At EVLP initiation, all grafts met Berlin criteria for ARDS based on the PaO_2_/FiO_2_ ratio^34^.

MSC treatment improved graft function during EVLP (Fig. 3a). The pulmonary vascular resistance was significantly lower in BM-MSC-treated (p=0.0247) and TAF-MSC-treated grafts (p=0.0057) than in non-treated grafts. The dynamic compliance was lower in non-treated grafts, although differences were not significant. PaO_2_/FiO_2_ was significantly higher in BM-MSC-treated (364.6±73.3; p=0.0427) and TAF-MSC-treated grafts (471.5±54.7; p=0.0003) than in non-treated lungs (198.3±93.1), with no difference between the two MSC-treated groups (p=0.1142). A PaO_2_/FiO_2_ ratio of at least 300 mmHg is required for clinical graft acceptance for transplantation. After four hours of EVLP, 10 of 12 BM-MSC-treated lungs and all TAF-MSC-treated lungs met this criterion, whereas none of the non-treated lungs did. These functional improvements were paralleled by improved gross morphology (Extended Data Fig. 2a) and reduced bronchial inflammation in MSC-treated compared with non-treated grafts (Extended Data Fig. 2b).

**Fig. 3:**
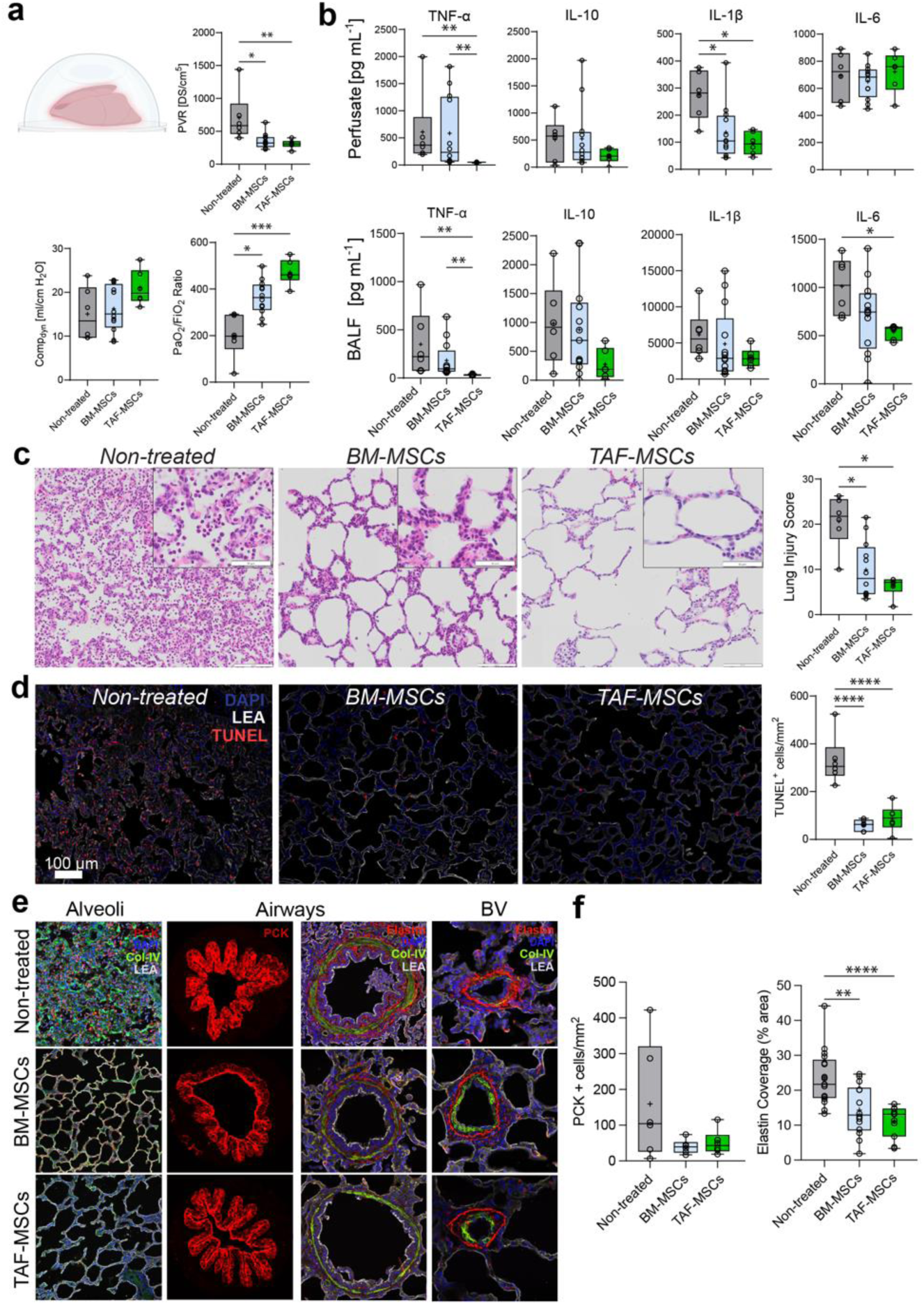
Improvement of graft function and inflammation following MSC treatment on ex vivo lung perfusion (EVLP). Following harvest and cold ischemic time, the donor lungs were connected to *ex vivo* lung perfusion for 4 hours. Non-treated grafts received placebo (n=6; grey), while treatment groups received a single dose of bone marrow–derived MSCs (BM-MSCs; n=12; light blue) or term amniotic fluid–derived MSCs (TAF-MSCs; n=6; green). All lungs were evaluated at the end of EVLP. **a)** Representative EVLP dome (created with Biorender.com) and clinically relevant parameters from hemodynamic monitoring and blood gas measurements, including pulmonary vascular resistance (PVR), dynamic compliance (Comp_dyn_), and ratio of partial pressure of arterial oxygen to fraction of inspired oxygen (PaO_2_/FiO_2_) after 4 hours on EVLP. **b)** Cytokine levels in EVLP perfusate (plasma, upper) and bronchoalveolar lavage fluid (BALF, lower). **c)** Representative haematoxylin and eosin (H&E) lung sections with scale bars representing 100 µm (overview) and 50 µm (callout). Quantification of histological lung injury scoring (right). **d)** Terminal deoxynucleotidyl transferase dUTP nick end labelling (TUNEL, red), 4′,6-diamidino-2-phenylindole (DAPI, blue) and Lycopersicon esculentum lectin (LEA, white) staining of lung tissue with quantification of TUNEL^+^ cells/mm^2^ (n=6, right). Scale bar represents 100 µm. **e)** Immunostaining for pan-cytokeratin (PCK, red) in alveoli (left column) and airways (second to left column), for elastin in airways (second to right column) and in blood vessels (right column) in non-treated upper), BM-MSC (middle) and TAF-MSC (lower row) groups. 4′,6-diamidino-2-phenylindole (DAPI, blue), Collagen IV (Col-IV, green) and Lycopersicon esculentum lectin (LEA, white) staining for visualization of tissue scaffold. **f)** Quantification of PCK^+^ cells/mm^2^ tissue (n=6 per group, left) and elastin coverage around blood vessels (% area) in different fields of view (n=17 non-treated, n=14 BM-MSCs, n=11 TAF-MSCs, right). Statistical comparisons were performed using one-way ANOVA or Kruskal-Wallis test when data were not normally distributed. *p<0.05, **p<0.01, ***p<0.001, ****p<0.0001. Boxplots: centre line, median; box, interquartile range; whiskers, min-max. If not otherwise stated, all animals were included in statistical analyses.

To assess inflammatory changes associated with functional recovery, cytokine concentrations were measured in BALF and perfusate (Fig. 3b). TAF-MSC treatment significantly reduced TNF-α (p=0.0031) and IL-1β (p=0.0239) in perfusate, and TNF-α (p=0.0016) and IL-6 (p=0.0258) in BALF. BM-MSC treatment significantly reduced IL-1β in perfusate compared with non-treated grafts (p=0.0350). TNF-α was also significantly lower in TAF-MSC-treated than in BM-MSC-treated grafts, both in perfusate (p=0.0073) and BALF (p=0.0060). No significant differences were observed in IL-10 or IL-6 in perfusate, or in IL-10 or IL-1β in BALF, across groups.

Histopathological analysis after EVLP showed significantly lower injury scores in BM-MSC-treated (p=0.0231) and TAF-MSC-treated grafts (p=0.0112) than in non-treated grafts, with no difference between the two MSC-treated groups (p>0.9999; Fig. 3c and Extended Data Fig. 2d). TUNEL staining showed reduced late apoptosis in both MSC-treated groups compared with non-treated grafts (p<0.0001 for both), with no difference between MSC-treated groups (p=0.7501; Fig. 3d). To assess structural remodelling, tissue sections were stained for PCK, elastin and SMA (Fig. 3e,f). Non-treated grafts showed greater epithelial disruption, whereas MSC-treated grafts showed preservation of vascular architecture, with significantly greater elastin coverage than non-treated grafts (p=0.005 and p<0.0001, respectively; Fig. 3e,f). During EVLP, white blood cell, lymphocyte and neutrophil counts showed the most pronounced differences between non-treated grafts and TAF-MSC-treated grafts (Extended Data Fig. 2c).

### Repeated dose MSC regimen sustained good lung function after lung transplantation

After four hours of EVLP, all lung allografts were transplanted into porcine recipients and closely monitored for three days (Supplementary Table 1; Fig. 4a). Recipients in the repeated-dose groups (n=6 BM-MSCs; n=6 TAF-MSCs) received two additional MSC infusions after transplantation, whereas the single-dose BM-MSC (n=6) and non-treated (n=6) groups received placebo. After three days of follow-up, right pneumonectomy rendered the animals fully dependent on the transplanted lung, and graft function was assessed for a further four hours.

**Fig. 4:**
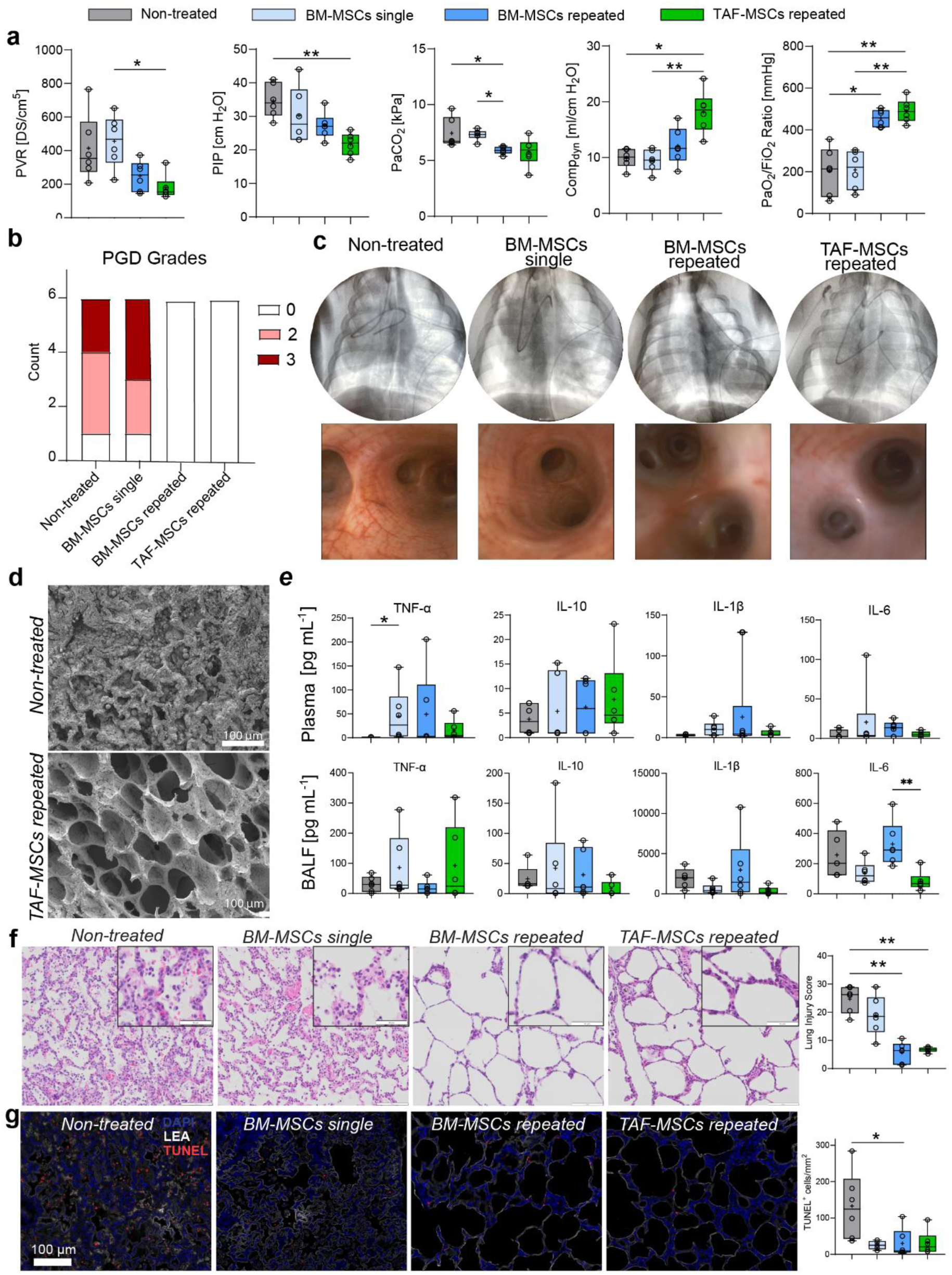
Repeated MSC treatment sustains lung function and reduces primary graft function after transplantation. The recipients were maintained under anaesthesia and monitored for three days after lung transplantation, after which a right pneumonectomy was performed to assess isolated graft function for 4 hours. Endpoints were compared across the four different groups (n=6): non-treated (grey), BM-MSCs single (light blue), BM-MSCs repeated (dark blue), and TAF-MSCs repeated (green). **a)** Clinically relevant parameters from hemodynamic monitoring and blood gas measurements, including pulmonary vascular resistance (PVR), peak inspiratory pressure (PIP), partial pressure of carbon dioxide (PaCO_2_), dynamic compliance (Comp_dyn_), ratio of partial pressure of arterial oxygen to fraction of inspired oxygen (PaO_2_/FiO_2_). **b)** Comparison of PGD grades at endpoint. **c)** Representative chest x-rays (upper) and bronchoscopy images (lower). **d)** Scanning electron microscopy (SEM) images of alveoli from non-treated (upper) and TAF-MSC-treated (lower) grafts. Scale bars: 100 µm. **e)** Cytokine levels in plasma (upper) and bronchoalveolar lavage fluid (BALF, lower). **f)** Representative haematoxylin and eosin (H&E) stained lung sections from the four different groups with scale bars representing 100 µm (overview) and 50 µm (callout). Quantification of histological lung injury scoring (right). **g)** Terminal deoxynucleotidyl transferase dUTP nick end labelling (TUNEL, red), 4′,6-diamidino-2-phenylindole (DAPI, blue), and Lycopersicon esculentum lectin (LEA, white) staining of lung tissue with quantification of TUNEL^+^ cells/mm^2^ (n=6, right). Scale bar represents 100 µm. Statistical differences were assessed by one-way ANOVA or Kruskal-Wallis test for not normally distributed data. *p<0.05, **p<0.01, ***p<0.001, ****p<0.0001. Boxplots: centre line, median; box, interquartile range; whiskers, min-max. If not otherwise stated, all animals were included in statistical analyses.

Repeated MSC dosing improved post-transplant graft function compared with the single-dose and non-treated groups (Fig. 4a). Repeated treatment reduced pulmonary vascular resistance (PVR), peak inspiratory pressure (PIP), and partial pressure of carbon dioxide (PaCO_2_), and improved dynamic compliance. In the repeated-dose TAF-MSC group, PIP was reduced (p=0.0039) and dynamic compliance improved (p=0.0132) relative to the non-treated group, whereas repeated-dose BM-MSC treatment reduced PaCO_2_ (p=0.0397). The PaO_2_/FiO_2_ was significantly higher in both repeated-dose groups than in the non-treated group, reaching 456.0±41.3 in the BM-MSC group (p=0.0423) and 491.2±56.4 in the TAF-MSC group (p=0.0049), compared with 202.7±114.1 in the non-treated group.

Chest X-ray evaluation together with PaO₂/FiO₂ ratios showed severe PGD (grades 2-3) in five of six animals in both the non-treated and single-dose BM-MSC groups, whereas no recipient treated with repeated MSC dosing developed PGD (Fig. 4b, c and Extended Data Fig. 3a). Scanning electron microscopy (SEM) analysis revealed marked differences in alveolar architecture between groups. Non-treated lungs displayed severe aspiration-induced injury with structural disruption and heterogeneous immune cell infiltration, whereas repeated MSC dosing preserved alveolar integrity and more closely resembled healthy morphology (Fig. 4d and Fig. 2d).

Cytokine profiling showed significantly higher plasma TNF-α in the single-dose BM-MSC group than in the non-treated group (p=0.0167). Repeated dosing of either MSC type did not differ from controls (Fig. 4e). In BALF, IL-6 was significantly lower in the repeated-dose TAF-MSC group than in the repeated-dose BM-MSC group (p=0.0046).

Histological analysis at the endpoint showed significantly lower lung injury scores in both repeated-dose MSC groups than in non-treated grafts (BM-MSCs, p=0.0060; TAF-MSCs, p=0.0069; Fig. 4f and Extended Data Fig. 3c). By contrast, single-dose BM-MSC treatment did not differ from non-treated grafts (p>0.9999). TUNEL staining showed the highest apoptosis rates in non-treated lungs, whereas all MSC-treated groups had fewer apoptotic cells, reaching significance in the repeated-dose BM-MSC group (p=0.0339; Fig. 4g).

### Repeated MSC dosing mitigated pro-inflammatory immune cell invasion

To assess the effects of MSC treatment on tissue morphology and pro-inflammatory immune cell invasion, endpoint sections were analysed by confocal imaging using PCK, elastin, and collagen IV (Fig. 5a). PCK quantification showed that non-treated and single-dose BM-MSC grafts retained levels comparable to those at initial lung injury, whereas repeated dosing with either MSC type reduced PCK signal to near-baseline values (Fig. 5b and Fig. 2g). Elastin staining indicated a trend towards consolidation in the repeated-dose groups (Fig. 5b).

**Fig. 5:**
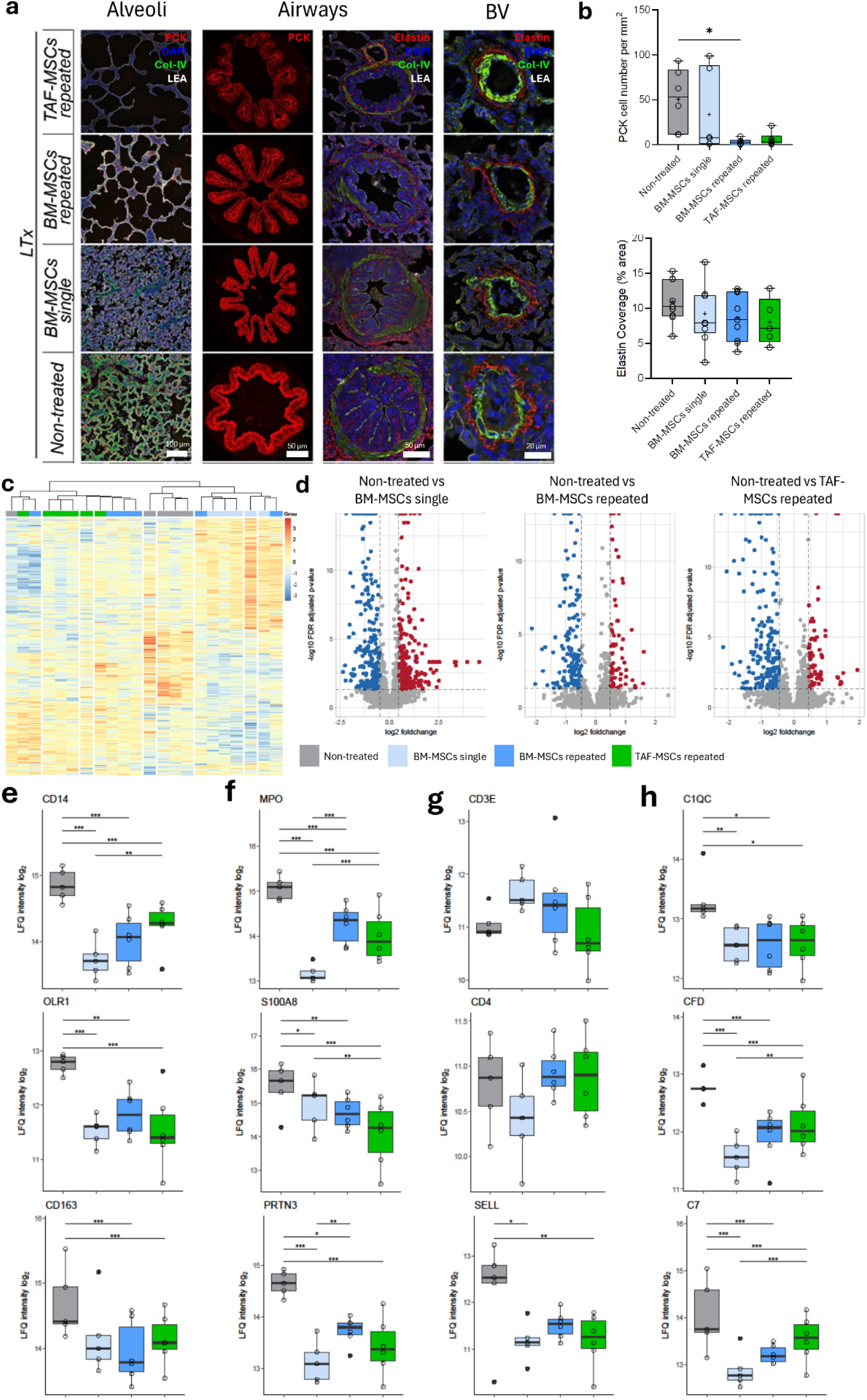
Repeated dose MSC treatment mitigates pro-inflammatory immune cell invasion. After three days monitoring and four hours isolated graft evaluation, tissue from all groups was analysed by confocal imaging and mass spectrometry (MS). **a)** Representative images of lung sections from the four different groups stained with pan-cytokeratin (PCK, red) in alveoli (left column) and airways (second to left column), and elastin staining in airways (second to right column) and in blood vessels (right column). 4′,6-diamidino-2-phenylindole (DAPI, blue), Collagen IV (Col-IV, green), and Lycopersicon esculentum Lectin (LEA, white) staining for visualization of tissue scaffold. Scale Bars: 100µm, 50µm, 50µm, 20µm in columns from left to right. **b)** Quantification of PCK^+^ cells/mm^2^ tissue (n=6, upper) and elastin coverage around blood vessels (% area) in different fields of view (lower, n=9 non-treated, BM-MSCs single, BM-MSCs repeated, n=5 TAF-MSCs repeated). **c)** Heatmap and hierarchical clustering of protein expression across individual samples from treated and non-treated groups. **d)** Volcano plots of differential protein expression comparing non-treated to TAF-MSCs (left), BM-MSC single (middle), and to BM-MSC repeated (right) with significantly overexpressed proteins in red and underexpressed in blue. **e-h)** Quantification of selected protein expression levels associated with monocytes **(e)**, neutrophils **(f),** lymphocytes (**g),** and the complement cascade (**h)**. All MS values are presented as log_2_ fold change in intensity, including Oxidized low density lipoprotein receptor (OLR1), Myeloperoxidase (MPO), S100 calcium binding protein A8 (S100A8), Proteinase 3 (PRTN3), CD3 epsilon chain (CD3E), L-Selectin (SELL), complement C1q C chain (C1QC), Complement Factor D (CFD), Complement C7 (C7). For **c)**-**h)** n=5 for non-treated, BM-MSCs single and n=6 for BM-MSCs and TAF-MSCs repeated. Statistical differences were assessed by one-way ANOVA or Kruskal–Wallis test for imaging data, and FDR-corrected p-values (q-values) for proteomic data. *p<0.05, **p<0.01, ***p<0.001. Boxplots: centre line, median; box, interquartile range; whiskers, min-max.

To determine whether these morphological differences were reflected at the proteomic level, we performed mass spectrometry on whole-tissue samples collected from all grafts at the endpoint. After filtering, 8,965 unique proteins were identified across groups, of which 1,227 were differentially expressed. Hierarchical clustering of these proteins showed clear separation of treatment groups, with non-treated grafts clustering distinctly from MSC-treated samples (Fig. 5c). Relative to non-treated grafts, 203 proteins were underexpressed and 68 overexpressed in TAF-MSC-treated lungs. In the single-dose BM-MSC group, 289 proteins were underexpressed and 279 overexpressed, whereas in the repeated-dose BM-MSC group, 148 proteins were underexpressed and 62 overexpressed (Fig. 5d). Across treatment groups, proteins associated with monocytes, neutrophils, lymphocytes, leukocyte rolling, and the classical and alternative complement pathways were consistently downregulated, while markers linked to tissue repair and regeneration were upregulated (Fig. 5e–h and Extended Data Fig. 4).

### Immune cell accumulation at the bronchial–vascular interface coincided with structural airway damage

Large field-of-view imaging of endpoint samples revealed pronounced elastin expression at the bronchial–vascular interface (BVI), the junction between airways and blood vessels (Extended Data Fig. 5a). Histological sections further identified the BVI as a site of increased immune cell infiltration in non-treated grafts at the endpoint (Extended Data Fig. 5b). To characterize this infiltrate in greater detail, we applied ultra-high-content imaging using the MACSima™ platform to endpoint graft samples from all four groups, representing the first use of this technology in a large-animal transplantation model. A total of 15 markers were imaged at subcellular resolution on the same tissue section, with approximately 1,068,960 cells individually segmented (Fig. 6a). CD3, CD4 and CD163 cell populations were increased in non-treated and single-dose BM-MSC grafts (Fig. 6b). Proliferation-associated markers, including histone H3, enhancer of zeste homologue 2 (EZH2) and Ki67, showed the same overall pattern (Extended Data Fig. 5c), consistent with the inflammatory and proliferative environment indicated by cytokine, histopathological and proteomic analyses. To examine airway–vascular remodelling in greater detail, we focused on the BVI in non-treated grafts (Fig. 6c and Extended Data Fig. 5d). High-resolution imaging revealed marked spatial polarization: CD163^+^ macrophages localized almost exclusively to the alveolar space, whereas CD3^+^ and proliferating cell nuclear antigen-positive (PCNA^+^) cells preferentially accumulated at bronchial and vascular sites, as confirmed by quantitative analysis (Fig. 6d).

**Fig. 6:**
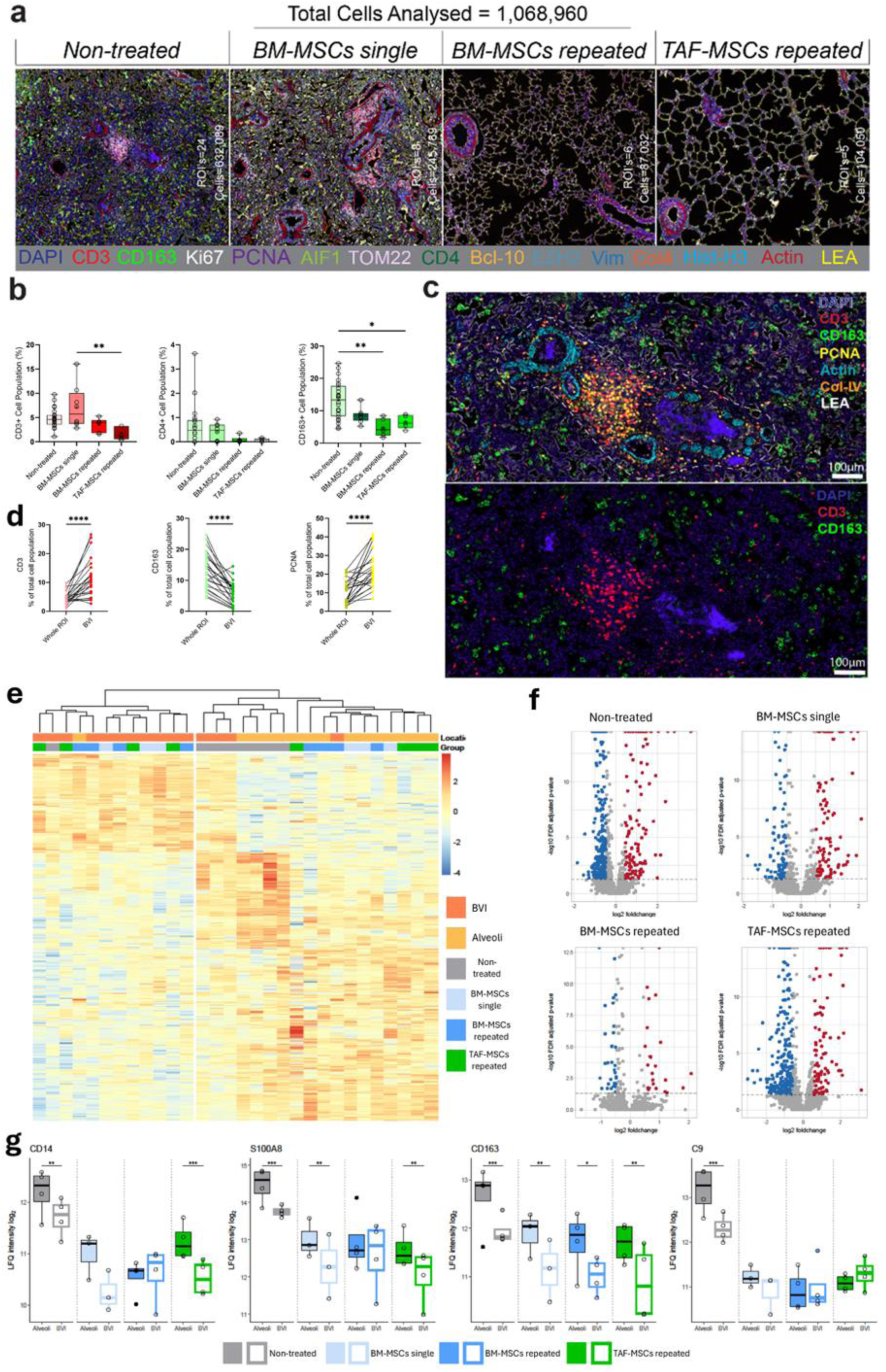
Immune cell invasion is enhanced at the bronchial-vascular interface (BVI). After three days monitoring and four hours isolated graft evaluation, tissue from all groups was analysed by 15-plex ultra-high content imaging with the (MACSima™) and by mass spectrometry following selective laser capture microdissection of regions of interest. **a)** 15-plex ultra-high content imaging of tissue from all groups at the endpoint. Total cells segmented and analysed = 1,068,960. **b)** Quantification of CD3^+^, CD4^+^, and CD163^+^ cell populations (% of whole population) in different regions of interest (ROI) in lung tissue at the end of experiment. (Non-treated n=23, BM-MSCs single n=8, BM-MSCs repeated n=6, TAF-MSCs repeated n=5). **c)** Representative images of bronchial-vascular interfaces (BVIs) from non-treated grafts at the endpoint. 4′,6-diamidino-2-phenylindole (DAPI), Proliferating cell nuclear antigen (PCNA), Collagen IV (Col-IV), Lycopersicon esculentum lectin (LEA). Scale bar: 100µm. **d**) Quantification of CD3^+^, CD163^+^, and PCNA^+^ cell populations comparing BVI and whole ROI across groups at the end of the experiment. **e)** Heatmap and hierarchical clustering of protein expression in alveoli (light orange) and bronchovascular interface (BVI, dark orange) across treated and non-treated groups. **f)** Volcano plots of differential protein expression comparing alveolar region and BVI within treatment groups. Significantly overexpressed proteins in red and underexpressed proteins in blue. **g)** Quantification of selected proteins associated with monocytes (CD14, CD163), neutrophils (S100 calcium binding protein A8 (S100A8)) and the complement system (Complement C9 (C9)); **e)**-**g)** n=4 per region and group for non-treated, BM-MSCs and TAF-MSCs repeated and n=3 for BM-MSCs single. Imaging data presented as mean ± SD. Statistical differences were assessed by using ANOVA or Kruskal-Wallis and Mann Whitney U-test for non-normally distributed data for imaging and FDR corrected p-values (q-values) for proteomic data. *p<0.05, **<0.01, ***p<0.001, ****p<0.0001. Boxplots: centre line, median; box, interquartile range; whiskers, min-max.

### The BVI emerged as a proteomic niche identified by laser capture microdissection

To further investigate the spatial differences suggested by advanced imaging, we compared alveolar regions and the bronchial–vascular interface (BVI) by mass spectrometry following laser capture microdissection. Alveolar and BVI areas were isolated from endpoint graft tissue sections across all groups. Unsupervised hierarchical clustering separated BVI from alveolar samples (Fig. 6e). In total, 5,261 unique proteins were identified. Of these, 384 were significantly differentially expressed between alveolar and BVI regions in the non-treated group, 192 in the BM-MSC single-dose group, and 55 in the BM-MSC repeated-dose group (Fig. 6f). Gene set enrichment analysis (GSEA) revealed regional differences across multiple biological processes, including wound healing, structural organization, immune and defence responses, cell communication, and responses to external stimuli (Extended Data Fig. 6a). In the non-treated group, these categories were broadly upregulated in alveolar relative to BVI samples, whereas fewer differences were observed in BM-MSC- and TAF-MSC-treated grafts, particularly after repeated dosing. Notably, when immune-related pathways were examined specifically, no pathways were differentially expressed in the BM-MSC repeated-dose group, in contrast to TAF-MSC-treated, BM-MSC single-dose, and non-treated grafts (Extended Data Fig. 6b). Protein-level analysis further highlighted region-specific signatures associated with aspiration-induced lung injury. Neutrophil-associated proteins, complement factors, macrophage markers, and structural proteins, with the exception of vimentin, were more highly expressed in alveolar regions than in the BVI (Fig. 6g and Extended Data Fig. 6c).

## Discussion

In this study, we show that repeated MSC dosing across EVLP and the early post-transplant period, but not single-dose treatment restricted to EVLP, provides durable rescue of severely aspiration-injured donor lungs. In a clinically relevant porcine lung transplantation model with 72-hour follow-up, repeated administration of both BM-MSCs and lung-specific TAF-MSCs restored graft function, reduced tissue injury, and prevented PGD. A single dose administered during EVLP improved early graft function but failed to sustain protection after transplantation. These findings identify dosing schedule, rather than cell source, as the key determinant of durable efficacy in this setting and establish repeated MSC delivery as a strategy to recover donor lungs that would otherwise remain non-transplantable.

Donor lung shortage remains a major limitation in lung transplantation and a principal cause of waitlist mortality. As many as 80% of donor lungs are ultimately discarded, with aspiration injury accounting for a substantial proportion of these cases^3,4^. Currently, there is no effective therapy for aspiration-induced ALI, which can progress to severe pneumonitis or ARDS, leading to exclusion of these grafts from transplantation^1,3^. In the present study, aspiration-induced ARDS was confirmed in all donor animals by the Berlin definition^34^, with impaired gas exchange and bilateral radiographic infiltrates, but also by a lack of right ventricular dysfunction, elevated inflammatory mediators, severe histopathological injury, and increased total bile acids^35–37^. EVLP provided a controlled platform for targeted delivery of MSCs^16,17,38^, and consistent with previous reports in ischemia-reperfusion injury and lipopolysaccharide-induced ARDS, a single intra-EVLP dose improved oxygenation compared to non-treated grafts^16,17,20^. However, unlike repeated treatment, this early benefit was insufficient to prevent post-transplant deterioration, indicating that aspiration-induced injury continues to evolve beyond the perfusion phase and requires sustained therapeutic modulation through reperfusion and the early post-transplant period^39^.

This distinction between transient rescue and durable protection is central. Although MSC treatment during EVLP restored function and rendered a substantial proportion of injured grafts suitable for transplantation, only repeated dosing prevented progression to PGD after transplantation and ensured successful graft survival. None of the recipients treated with repeated MSC dosing developed PGD, whereas five of six recipients in both the untreated and single-dose BM-MSC groups developed the most severe PGD grade 3. These findings suggest that MSC therapy must be aligned not only with the ex vivo perfusion phase but also with the subsequent inflammatory stresses of implantation and reperfusion. More broadly, they support a treatment paradigm in which regenerative cell therapy is initiated during EVLP and continued through the early post-transplant period, rather than being restricted to organ preservation alone^16^.

Both BM-MSCs and TAF-MSCs were effective under repeated administration. While TAF-MSCs showed a non-significant trend towards greater benefit in some readouts, the central finding is that treatment efficacy was driven primarily by the dosing regimen rather than cell source, as both MSC types provided comparable protection when given repeatedly. This is important for translation, as it shifts the focus from identifying a single optimal MSC source to designing therapeutically effective dosing strategies that can be implemented within existing EVLP and transplantation workflows. We previously showed that TAF-MSCs, administered under the same repeated regimen, restored function in lipopolysaccharide-injured donor lungs and prevented PGD^20^. The present findings extend this evidence to aspiration injury and indicate that, in severely injured donor lungs, regimen design is more decisive than tissue origin for durable graft protection. This is particularly relevant given that previous *in vitro* studies have reported source-dependent differences in MSC immunomodulatory potency and proliferative capacity^24–26^, but direct *in vivo* comparisons remain scarce.

The multimodal analyses in this study further suggest that repeated MSC dosing broadly attenuates early inflammatory injury while supporting structural recovery. Proteomic analyses showed downregulation of proteins associated with monocytes, neutrophils, lymphocytes, leukocyte rolling, and complement activation in treated grafts, together with upregulation of markers linked to tissue repair and regeneration. Within the monocyte/macrophage compartment, inflammatory markers were reduced across treated groups, consistent with an overall attenuation of myeloid recruitment and activation. Although CD163-positive cells were also reduced at the endpoint, in contrast to reports suggesting MSC-supported M2 polarization^22^ this most likely reflects reduced macrophage infiltration overall rather than failure to support reparative polarization. Consistent with this regenerative profile, Nestin was upregulated across treatment groups, supporting the notion that MSC therapy not only dampened immune activation but also promoted tissue recovery^40^.

Neutrophil-associated injury appeared to be a particularly prominent component of this response. Repeated MSC treatment reduced circulating neutrophil activity and decreased tissue abundance of proteins associated with neutrophil effector function and neutrophil extracellular trap (NET) formation, including azurocidin 1 (AZU1), myeloperoxidase (MPO), S100 calcium-binding protein A8 and A9 (S100A8 and S100A9), and proteinase 3 (PRTN3). NETs are central mediators in the pathogenesis of ALI^15,37,41^ and are increasingly recognized as contributors to graft dysfunction after lung transplantation. Their suppression therefore likely constitutes a key mechanism of MSC-mediated graft protection. This is supported by recent studies showing that direct NET removal during EVLP improved function and morphology in aspiration-damaged donor lungs^15^, while cytokine adsorption reduced circulating nucleosome levels, inflammatory injury, and primary graft dysfunction in experimental and clinical lung transplantation settings^14,42^. This observation also aligns with our previous study in LPS-injured lungs, where TAF-MSCs similarly reduced proteins associated with neutrophil degranulation and aggregation during EVLP and post-transplantation^20^.

Complement inhibition may represent an additional mechanism. Treated grafts showed reduced abundance of proteins linked to the classical pathway (C1QB, C1QC), the alternative pathway (CFD), and the terminal membrane attack complex (C5-C9)^43^. Because complement activation has been implicated in PGD in both preclinical and clinical transplantation^44–46^, coordinated downregulation of complement-related proteins further supports the idea that repeated MSC dosing interrupts multiple interconnected inflammatory cascades that drive early graft dysfunction.

A notable finding of the spatial analyses was the identification of the bronchial-vascular interface (BVI) as a focal site of immune activation in aspiration-damaged lungs. High-content imaging revealed preferential accumulation of CD3-positive and proliferating cell nuclear antigen-positive cells at the BVI, whereas CD163-positive macrophages localized predominantly to the alveolar space, indicating compartment-specific inflammatory organization in untreated grafts. Anatomically, the BVI lies within peribronchovascular and perivascular regions that have been implicated in early oedema and leukocyte infiltration^47,48^. The prominence of CD3-positive cells in untreated grafts and their reduction in MSC-treated groups are consistent with MSC immunomodulatory effects and suggest that T-cell modulation may contribute to the therapeutic response^26^. Proteomic analyses after laser capture microdissection confirmed that the BVI represents a distinct molecular niche. Untreated and single-dose grafts retained marked regional differences between alveolar and BVI compartments, whereas repeated MSC therapy attenuated these differences and produced a more homogeneous immune landscape. These findings suggest that repeated MSC dosing not only reduces overall inflammation but also normalizes its spatial distribution within the graft. Taken together, the data identify the BVI as a critical site of early immune activation in aspiration-damaged lungs and suggest that restoring immune balance between alveolar and bronchial-vascular compartments may be one mechanism by which repeated MSC therapy mitigates early PGD pathogenesis.

From a translational perspective, both human BM-MSCs and TAF-MSCs were well tolerated, with no observed adverse effects. Although BM-MSC expansion was performed outside a GMP facility, production and quality control were carried out under GMP-compatible conditions, supporting feasibility for clinical development. MSCs have already been explored in early trials for ARDS and lung transplantation, but repeated peri-transplant dosing and direct comparison of distinct MSC sources have not previously been investigated in this context^29–31,49^. Our findings therefore provide a rationale for integrating repeated MSC administration into EVLP-based rehabilitation protocols for severely injured donor lungs.

This study was designed to evaluate early graft recovery in a clinically relevant porcine aspiration model and therefore focused on the first 72 hours after transplantation, the period in which PGD develops and early therapeutic effects are most critical. Although longer-term outcomes, additional donor-lung injury phenotypes, and other cell therapy products were beyond the scope of the present work, the model captures a severe and clinically important form of donor lung injury. Likewise, while the mechanistic analyses were not designed to establish direct causal contributions of individual pathways, the convergence of physiological, histological, spatial and proteomic findings provides a robust framework for understanding the therapeutic effects of repeated MSC administration.

In summary, repeated MSC administration across EVLP and the early post-transplant period was required to sustain successful graft function, reduce immune-mediated injury, and prevent PGD in severely aspiration-injured donor lungs, whereas MSC treatment limited to EVLP restored graft function sufficiently for transplantation, it did not provide durable protection after transplantation. Both BM-MSCs and TAF-MSCs were effective under repeated dosing, identifying treatment schedule rather than cell source as the principal determinant of efficacy. These findings support repeated perfusion-guided MSC therapy as a clinically compatible regenerative strategy for restoration of severely injured donor lungs and support a broader shift from donor-organ triage towards donor-organ repair. More broadly, they suggest that combining machine perfusion with rationally designed peri-transplant cell therapy may provide a generalizable framework for organ rescue across solid-organ transplantation.

## Methods

Refer to Supplementary Methods for greater detail of subsections.

### Ethical Considerations and Animal Preparation

All animals were given care according to the USA principles of Laboratory Animal Care of the National Society for Medical Research, Guide for the Care and Use of Laboratory Animals, National Academies Press (1996). All animal handling, welfare monitoring, and euthanasia were performed according to the guide and under the supervision of an on-site veterinarian and approved by the local Ethics Committee for Animal Research (Dnr 5.2.18-4903/16, and Dnr 5.2.18-8927/16) at Lund University.

In total, 48 adult Yorkshire pigs, 24 donors and 24 recipients with a mean weight of 40 kg, were included in the study. Paired donors and recipients were randomized into four experimental groups: pigs treated with BM-MSCs during EVLP only (BM-MSCs single), pigs treated with BM-MSCs during EVLP and post transplantation (BM-MSCs repeated), pigs treated with lung-specific term amniotic fluid-derived MSCs (TAF-MSCs repeated), and pigs receiving no MSC treatment at all (non-treated). A single-dose TAF-MSC group was not included because the study was powered to test the effect of repeated dosing and to compare MSC sources under the repeated-dose regimen, while minimizing animal use. An overview of the experimental timeline and sampling scheme are presented in Fig. 1. All pigs received premedication and general anaesthesia as previously reported^14,20^.

### Induction and establishment of aspiration lung injury

Standardized gastric content was used to induce lung injury in all donor pigs. Gastric contents had been previously retrieved from pigs, aliquoted, and stored at −80°C. Prior to the experiment, the gastric content was thawed, homogenized, centrifuged, and filtered to remove all particulates and finally titrated to a pH of 2.0 before instillation. Each donor pig received 2 mL/kg/lung (in total 4 mL/kg) equally distributed throughout the lung lobes via bronchoscopy, given over two doses. The first consisted of 90% of the total dose of gastric content, and the second one of the remaining 10% was administered one hour later. Subsequently, the donor pigs remained under anaesthesia for 6 hours for the establishment of the lung injury. A chest x-ray was taken before the start of induction of lung injury, and hourly thereafter. Induction of acute lung injury was measured by PaO_2_/FiO_2_ and further confirmed retrospectively through histological assessment^34^.

### Monitoring through hemodynamic and arterial blood gases

Arterial blood gases were analysed every 30 minutes in the donor animals and during the last four hours of recipient follow-up. During EVLP and the first 60 hours of recipient follow-up, blood gases were drawn hourly and analysed with an ABL 90 FLEX gas analyser (Radiometer Medical ApS, Brønshøj, Denmark). Haemodynamic parameters were measured every 30 minutes in the donor animals and then hourly during EVLP and the first 60 hours of recipient follow-up. In the last four hours of recipient follow-up, the hemodynamic parameters were measured every 30 minutes.

### Pulmonary harvest after established lung injury - Donor

After lung injury was established in the donors, a median sternotomy was performed. With the use of a 28 F cannula secured by a purse-string suture, the pulmonary artery was cannulated via the right ventricle and placed in the outflow tract of the pulmonary artery. A clamp was put on the inferior vena cava, the superior vena cava, and the ascending aorta whereas the inferior vena cava and the left atrium were left open. Subsequently, lungs were perfused at a perfusion pressure of < 20 mmHg with distribution of 4 L of cold Perfadex^®^ solution (XVIVO perfusion, Gothenburg, Sweden) using anterograde perfusion. After four hours of perfusion, lungs were harvested *en bloc* using standard procedure. After harvest, lungs were stored at 4°C for two hours, fully immersed in Perfadex^®^ solution.

### Ex vivo lung perfusion (EVLP)

EVLP was performed using a Vivoline LS1 platform (XVIVO perfusion, Gothenburg, Sweden). Target perfusion was set at 40% of cardiac output, a tidal volume of 7 mL/kg body weight of the donor, PEEP of 5 cm H_2_O, respiratory rate (RR) of 7 FiO_2_ at 21% for four hours^50^. The system was primed with red blood cells drawn from the donor animal to achieve a haematocrit level of 15-20% in the EVLP circuit, as well as with Steen^TM^ Solution (XVIVO perfusion, Gothenburg, Sweden).

### Treatment with bone-marrow-derived and full-term amniotic fluid-derived MSCs

Human bone marrow (BM) was collected from young healthy consenting volunteers. BM was aspirated as approved by the Swedish Ethical Review Authority at the Haematology Department, Lund, Sweden. Cells were isolated and propagated using a GMP-compatible animal serum-free culturing protocol using pooled human platelet lysate as described previously and stored frozen in early passage until use^51^. The cultured MSCs were characterized by surface marker expression via flow cytometry and differentiation assays, and tested for their immunomodulatory potential^52^.

TAF-MSCs, as previously described, were obtained and isolated from the amniotic fluid of healthy donors who provided informed consent and were undergoing scheduled full-term caesarean section deliveries^20^. The subsequent manufacturing and quality control were in accordance with good manufacturing practice. For this study, the cells were kept frozen at −150°C until one hour before use. After thawing, all MSCs used in this study were washed (centrifuged for 5 min at 350xg and then resuspended in 50 mL PBS). Cell counts and viability were determined with a NucleoCounter NC-250 (Chemometec, Allerød, Denmark).

The treated groups received either BM-MSCs or TAF-MSCs at a dose of 2 × 10^6^ cells per kg body weight at one or three time points: during EVLP (BM-MSC single dose) and then 1 hour and 12 hours after transplantation (BM-MSCs repeated dose, TAF-MSCs repeated dose). The non-treated group received a placebo treatment in the form of PBS at the same timepoints, all administered intravenously. Each dose was given over a 20-30-minute timeframe to avoid increasing PVR^35^.

### Left lung transplant recipient

The lung transplantation was performed according to the protocol described by Mariscal *et al.*^53^. In brief, a left thoracotomy was performed, and the pulmonary artery, atrium, and bronchus were clamped sequentially before pneumonectomy of the native lung. Donor grafts were implanted with bronchial anastomosis using polydioxanone sutures (PDS 4-0, Ethicon, Somerville, NJ, USA) and continuous atrial and arterial anastomoses with polypropylene sutures (Prolene 5-0, Ethicon). Recipients received tacrolimus (0.15 mg/kg, orally; Sandoz AS, Copenhagen, Denmark) and methylprednisolone sodium succinate (1 mg/kg, intravenously; Solumedrol, Pfizer, New York, NY, USA), and an open bronchial anastomosis was confirmed by bronchoscopy.

### Recipient follow-up

Post-transplantation, the animals were kept under anaesthesia for three days as previously described^14,20^. In brief, anaesthesia was maintained with continuous infusions of ketamine (Ketaminol® vet, Intervet AB, Stockholm, Sweden), midazolam (Midazolam Panpharma®, Panpharma Nordic, Oslo, Norway), fentanyl (Leptanal®, Piramal Critical Care B.V., Lilly, France), and rocuronium bromide (Esmeron®, Merck, Kenilworth, NJ, USA). Imipenem (500 mg; Merck & Co. Inc., Kenilworth, NJ, USA) was administered intravenously three times daily, and dihydrostreptomycinsulfat (0.1 mL/kg; Boehringer Ingelheim Animal Health Nordics A/S) was given subcutaneously once per day. Immunosuppression was maintained with tacrolimus (0.15 mg/kg, orally; Sandoz AS, Copenhagen, Denmark) and methylprednisolone sodium succinate (1 mg/kg, intravenously, twice daily; Solumedrol, Pfizer, New York, NY, USA). Mechanical ventilation was adjusted to maintain adequate oxygenation while minimizing airway pressures, with PEEP set at 5–10 cmH₂O and peak inspiratory pressure kept below 30 cmH₂O.

### Right pneumonectomy

After three days of follow-up, the pulmonary hilum was dissected through a mid-sternotomy. To assess the isolated function of the transplanted left lung, a right pneumonectomy was performed, followed by an additional four hours of monitoring before the experiment was terminated, representing 68-72 hours of follow-up. The recipient was monitored using a Swan-Ganz catheter as described previously.

### Primary graft dysfunction determination and staging

Primary graft dysfunction (PGD) was determined by PaO_2_/FiO_2_ ratio and presence of infiltrates on imaging according to the ISHLT’s guidelines^54^. PGD grade 0 requires the absence of pulmonary oedema on the thoracic imaging and a PaO_2_/FiO_2_ ratio of at least 300 mmHg. Pulmonary oedema on imaging results in PGD grades 1-3 which are then distinguished by the degree of hypoxemia based on the PaO_2_/FiO_2_ ratio. Grade 1 is defined by a PaO_2_/FiO_2_ ratio lower than 300 mmHg, Grade 2 between 200-300 mmHg, and grade 3 at 200 mmHg or less. Presence of lung infiltrates was assessed through imaging with a mobile C-arm x-ray machine (Siemens, Munich, Germany).

### Histopathological analysis

Lung biopsies were obtained from the lungs of healthy pigs (baseline), after confirmation of lung injury, at the start and end of EVLP, and at termination of the experiment. Fresh tissue was fixed in 10% Formalin, processed, embedded, and sections cut according to standard histopathological protocol. Dried sections were stained with Hematoxylin& Eosin according to protocol and imaged using a VS-120 slide scanner (Olympus, Tokyo, Japan).

### Molecular measurements

Cytokine analysis was performed on plasma, EVLP perfusate, and bronchoalveolar lavage fluid (BALF) using the Cytokine & Chemokine 9-Plex Porcine ProcartaPlex™ Panel 1 (Thermo Fisher Cat. No. EPX090-60829-901, Thermo Fisher Scientific, Waltham, Massachusetts, US) kit according to manufacturer’s instructions.

Amount of total bile acids (TBA) was analysed in BALF using a Bile Acid Assay Kit (ab239702, Abcam, Cambridge, United Kingdom) according to manufacturer’s instructions.

### Differential blood cell counts

Differential counts of lymphocytes, neutrophils, and total white blood cell counts were determined in EDTA anti-coagulated whole blood (BD Vacutainer, BD Biosciences) using a Sysmex KX-21N automated haematology analyser (Sysmex, Milton Keynes, UK).

### Confocal Tissue Processing and Imaging

Ten µm FFPE lung sections were de-paraffinized and rehydrated according to standard protocols. After antigen retrieval and staining, images were acquired using a Nikon A1RHD equipped with a 20x objective (NA-0.8) and piezo 200 stage.

TUNEL Assay Kit BrdU Red was used (Abcam, ab66110) according to suppliers’ instructions and samples imaged on a Nikon Ti2 epifluorescence microscope (Nikon, Amstelveen, Netherlands).

### SEM Tissue Processing and Imaging

Lung tissue samples were embedded in 3% low melting agarose (Sigma-Aldrich, Darmstadt, Germany) and cut into 200 µm sections using a vibrating microtome (Leica VT1200S, Leica, IL, US). Samples were subjected to consecutive washes, fixation, and drying steps and imaged on a Jeol JSM-7800F FEG SEM (JEOL, Tokyo, Japan).

### Ultra-High-Content Tissue Processing and Imaging

Ten µm FFPE lung sections were de-paraffinized and rehydrated according to standard protocols. Antigen retrieval was carried out in TEC buffer for 30min at 98°C. Imaging was carried out in the MACSima™ platform (Miltenyi Biotec, Bergisch Gladbach, Germany) and prepared as previously described^55^.

### Proteomic analysis with mass spectrometry

Homogenized lung tissue was processed for protein extraction and enzymatic digestion using S-Trap protocols, and peptides were analyzed on a TIMS TOF HT mass spectrometer with data-independent acquisition as previously described^20^. Raw spectra were processed with DIA-NN v1.8.1, and normalization and differential expression analyses were performed in R (v4.3.1) using MS-DAP^56^ with MS-EmpiRe^57^. Significantly differentially expressed proteins were defined as FDR <0.05 and bootstrapped log2 fold-change cutoff values.

Laser-capture microdissection was performed on alveolar regions and the bronchial–vascular interface, and collected material was processed and analyzed by mass spectrometry using the same workflow. Functional annotation and enrichment for both datasets were performed with PANTHER GO analysis.

### Statistical analysis

Data are reported as mean ± standard deviation (SD) and presented as boxplots with individual values, median lines, a plus (+) at the mean, and whiskers indicating minimum and maximum in their graphical form. Normal distribution of the data was tested with Shapiro-Wilks test.

Statistical differences between and within groups were tested with Student’s t-test when data were normally distributed. When data were not normally distributed, nonparametric tests were used including the Wilcoxon test within groups and the Mann-Whitney U-test between groups. One-way ANOVA was used within groups when the data were normally distributed and the Friedman test when it was not normally distributed for paired data or Kruskal Wallis test for non-paired data. Observed frequencies of categorical variables were compared with a Chi-squared test. The mixed effects model with Šídák’s correction for multiple comparisons was used with repeated measures data. Statistical analyses were performed with GraphPad Prism (version 10.1, GraphPad Software, San Diego, USA) or R Studio (version 4.2.2), and significance was defined as *p<0.05, **p<0.01, ***p<0.001, ****p<0.0001.

## Acknowledgements

We express our deep gratitude for perfusionist Dr Leif Pierre for contributing to this research with his tremendous expertise in EVLP and porcine experiments to establishing and performing these experiments. Without him, this project would have not been possible. The authors acknowledge Amniotics AB, for donating the cells used in this study, and Sven Kjellström at BioMS, Lund University, Lund, Sweden for support with the MS.

## Funding

The authors gratefully acknowledge funding received by the Swedish research council (SL), the EU Interreg Öresund-Kattegat-Skagerrak (SL), by grants from the Swedish state under the agreement between the Swedish government and the county councils, and by the ALF-agreement, (SL).

## Author Contributions

SL and FO designed the study. SL, MM, NBB, DE, AN, MS, EB, GH, HG, GK, SH, and FO performed the animal experiments and collected measurements or samples during the animal experiments. FO, MM, AN, NBB, GH, and SL acquired the data and performed data analysis. SS provided access to and aspirated the BM and provided MSC expertise. MO and KS provided critical protocols and materials for GMP compatible MSC expansion and expertise. FO, MM, NBB, AN, and SL prepared the manuscript. All authors have read and approved the final version.

## Competing interests

S.L. has received a state research grant (Vinnova) with Amniotics AB, Lund, Sweden, as a co-applicant. All other authors declare no conflicts of interest.

## Extended Figures and extended Figure legends

**Extended Data Fig. 1:**
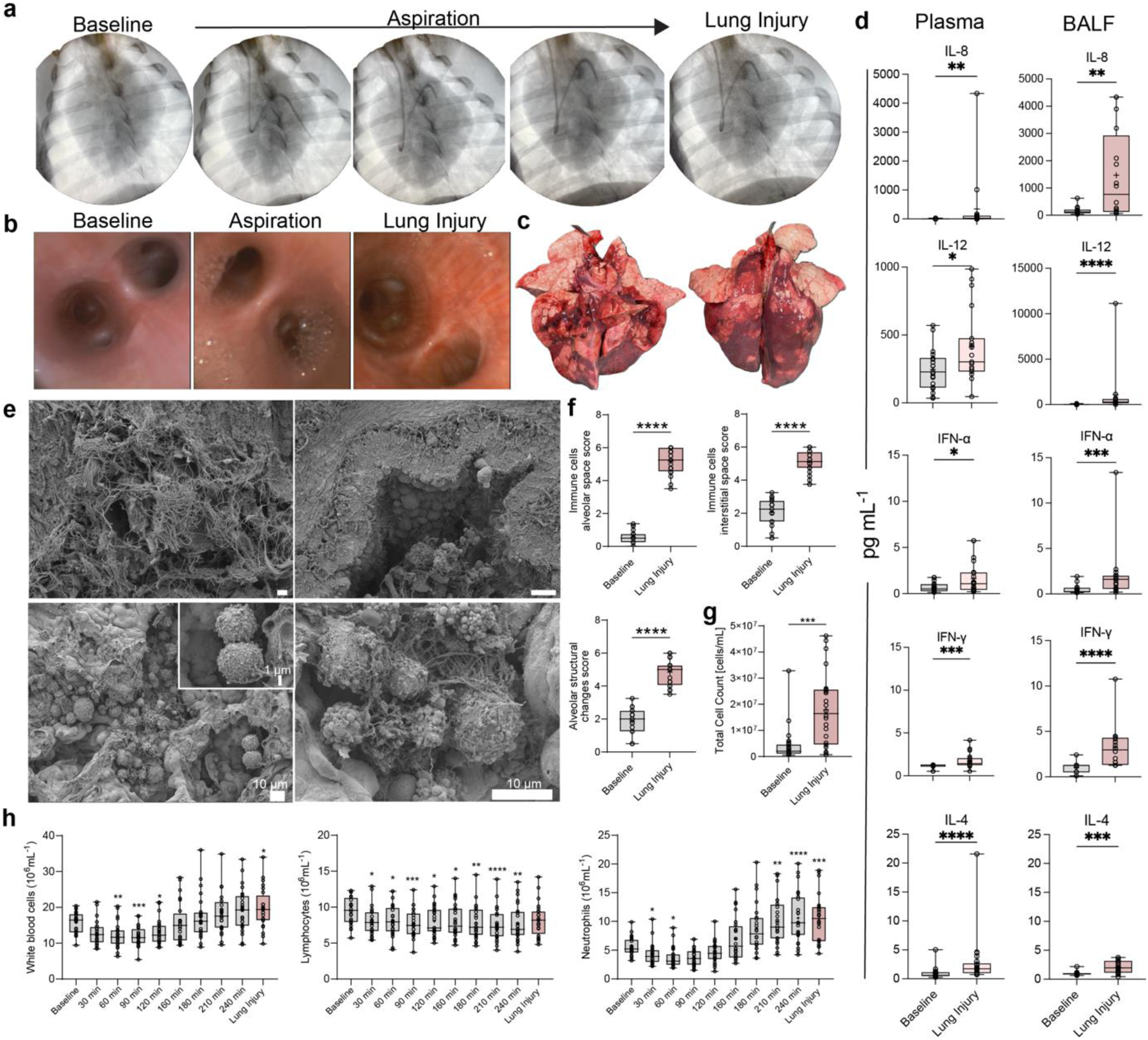
Establishment of aspiration-induced lung injury. Healthy animals (Baseline) received gastric contents (aspiration) to induce acute lung injury (lung injury). **a)** Chest x-rays taken over the course of lung injury induction. **b)** Bronchoscopy images captured at baseline, during aspiration, and after confirmation of lung injury. **c)** Lung gross morphology at lung injury. **d)** Cytokine levels at baseline and after lung injury in plasma (n=17 and n=22, respectively, left column) and bronchoalveolar lavage fluid (BALF, n=16 per group, right column). **e)** SEM images of damaged cilia in bronchi (upper panel) and immune cell clusters (lower panel) in alveolae after confirmation of lung injury. Scale bars represent 10 µm and 1 µm (callout). **f)** Subcategories of lung injury scoring comparing baseline to lung injury: immune cells in alveolar spaces, immune cells in interstitial spaces and alveolar structural changes. **g)** Total cell counts (cells/mL) in BALF at baseline and after confirmation of lung injury. **h)** Amount of white blood cells, lymphocytes, and neutrophils measured in whole blood over the course of injury induction. Statistical differences were calculated by using ANOVA or Kruskal-Wallis test when data were not normally distributed, and Mann-Whitney U-test. 05, **<0.01, ***p<0.001, ****p<0.0001. Boxplots: centre line, median; box, interquartile range; whiskers, min-max. If not otherwise stated, all animals were included in statistical analyses.

**Extended Data Fig. 2:**
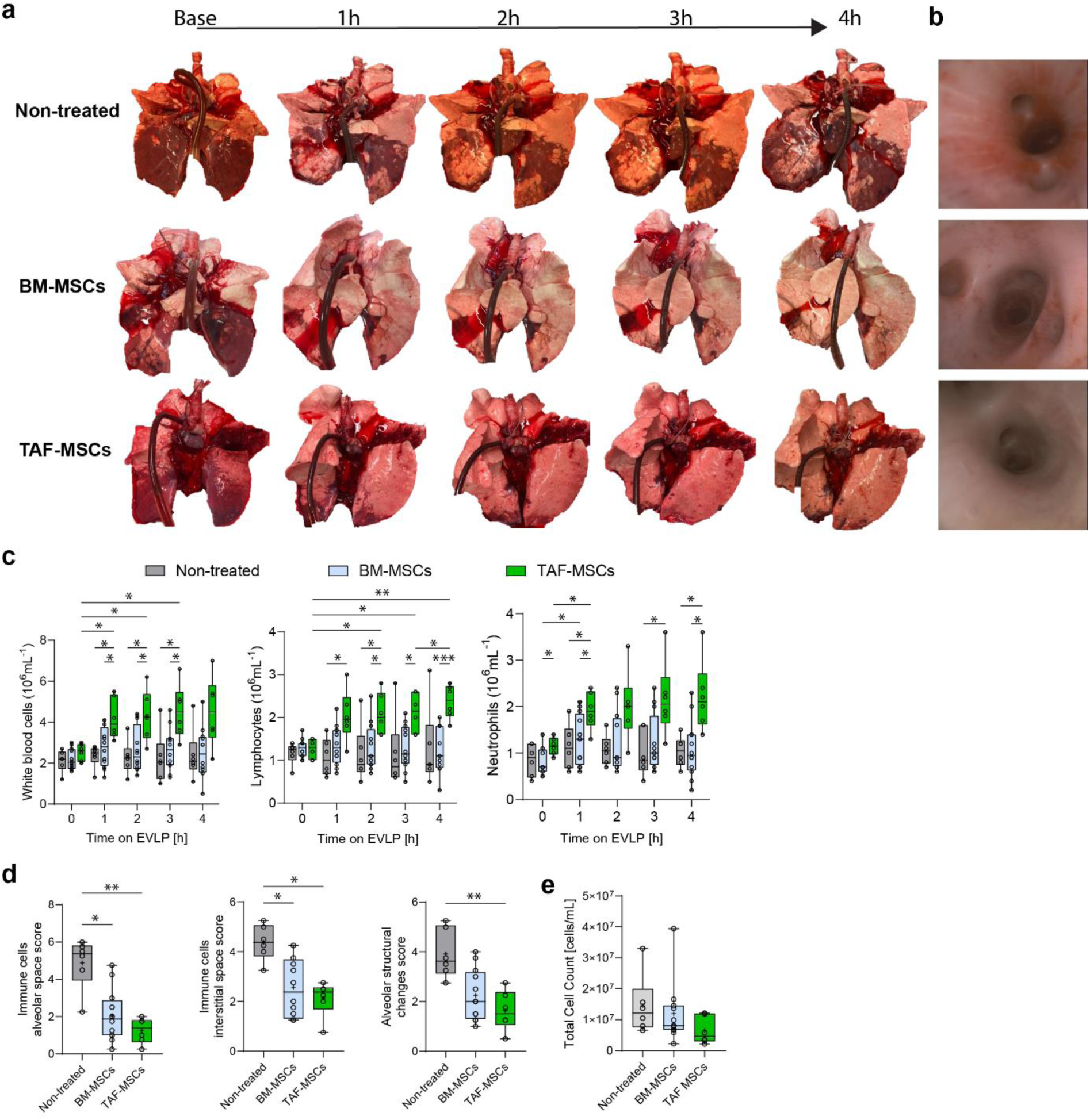
Morphological and cellular status improve during *ex vivo* lung perfusion (EVLP) with MSC treatment. Lungs with confirmed lung injury were explanted and placed on EVLP for 4 hours. The three groups received EVLP with one dose of either placebo (non-treated, grey; n=6), bone marrow-derived mesenchymal stromal cell (BM-MSCs, light blue; n=12), or term amniotic fluid-derived mesenchymal stromal cells (TAF-MSCs, green; n=6). **a)** Representative gross morphology of lungs during 4 hours of EVLP. **b)** Representative bronchoscopy images at the end of EVLP. **c)** White blood cell, lymphocyte, and neutrophil counts in EVLP perfusate during EVLP. **d)** Subcategories of lung injury scoring after 4 hours of EVLP: immune cells in alveolar spaces, immune cells in interstitial spaces, and alveolar structural changes. **e)** Total cell count (cells/mL) in bronchoalveolar lavage fluid. Statistical differences were calculated by using ANOVA or Kruskal-Wallis test when data were not normally distributed. *p<0.05, **p<0.01, ***p<0.001, ****p<0.0001. Boxplots: centre line, median; box, interquartile range; whiskers, min-max. If not otherwise stated, all animals were included in statistical analyses.

**Extended Data Fig. 3:**
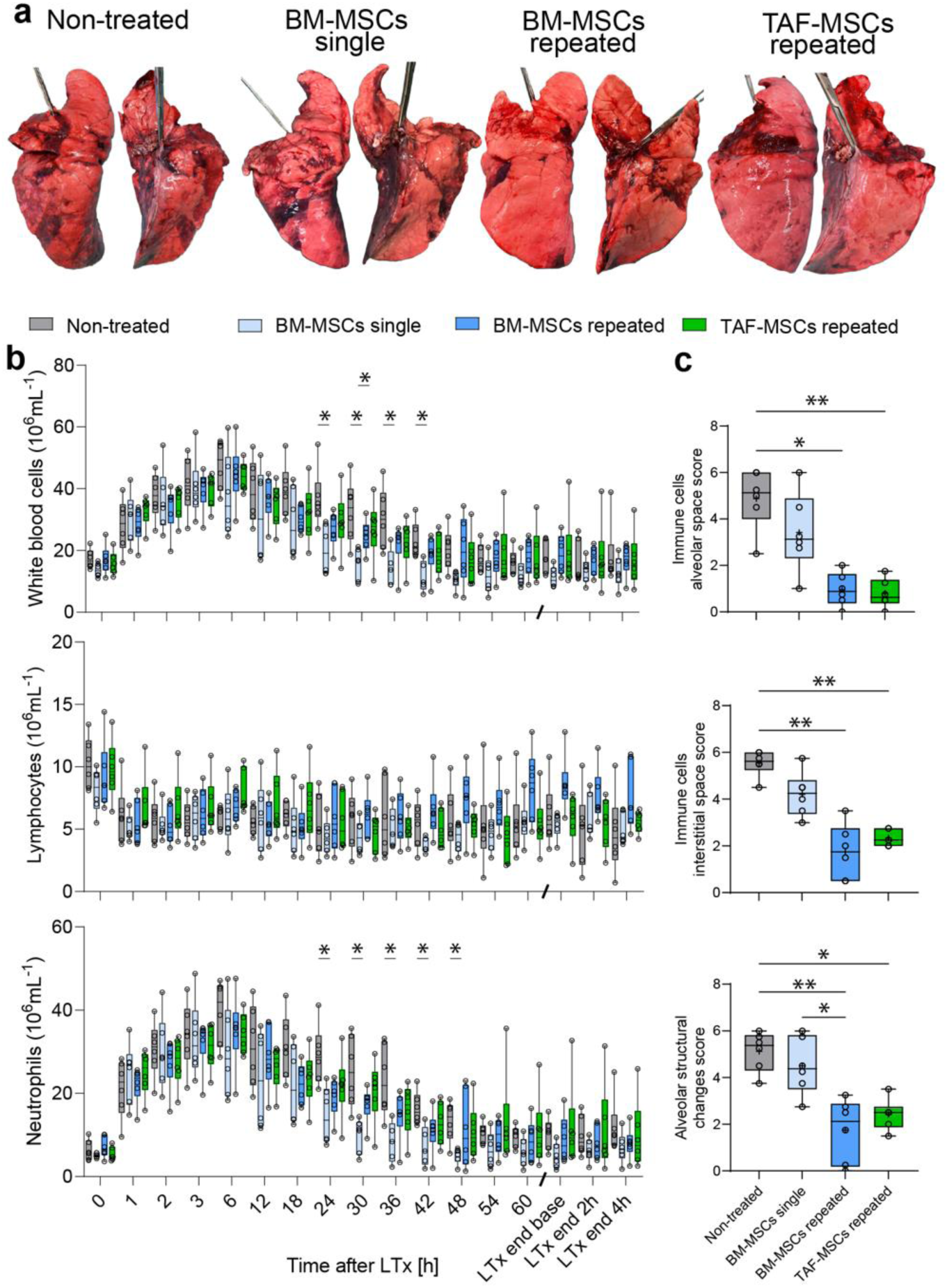
Improved macroscopic and cellular status is sustained post-transplantation following treatment with repeated doses of MSCs. After left lung transplantation, recipients were monitored for three days, followed by 4 hours of isolated evaluation of the left lung. All 4 groups are compared: non-treated (n=6, light grey), BM-MSCs single (n=6, light blue), BM-MSCs repeated (n=6, dark blue), and TAF-MSCs repeated (n=6, green). **a)** Representative gross morphology of the explanted left lungs at endpoint, front and back. **b)** White blood cell, lymphocyte, and neutrophil counts following transplantation and 4-hours of isolated lung function assessment (LTx end). **c)** Subcategories of lung injury scoring at the endpoint: immune cells in alveolar spaces, immune cells in interstitial spaces, and alveolar structural changes. Statistical differences were calculated by using ANOVA or Kruskal-Wallis test when data was not normally distributed. *p<0.05, **p<0.01, ***p<0.001, ****p<0.0001. Boxplots: centre line, median; box, interquartile range; whiskers, min-max. If not otherwise stated, all animals were included in statistical analyses.

**Extended Data Fig. 4:**
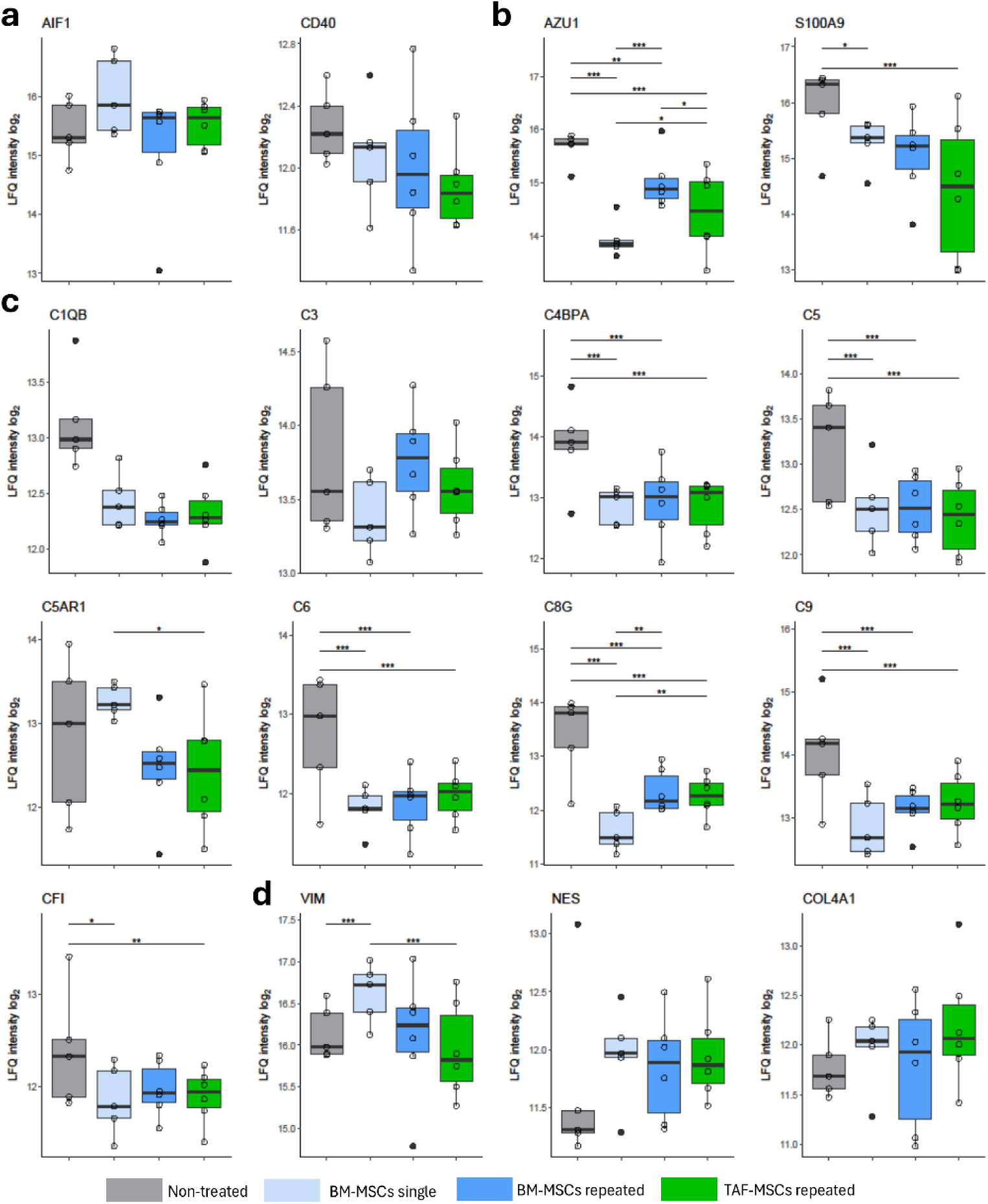
Repeated dose MSC treatment quells proteomic pro-inflammatory signature post transplantation. After three days of monitoring and four hours of isolated graft evaluation, tissue from all groups was analysed using mass spectrometry. **a-c)** Quantification of selected protein expression levels associated with monocytes and lymphocytes **a)**, neutrophils **b),** the complement cascade **c),** and structural integrity **d),** see Supplementary Table 2 for protein abbreviations. All individual values are presented as log_2_ fold change in intensity with FDR corrected p-values (q-values): *p<0.05, **p<0.01, ***p<0.001, ****p<0.0001. Boxplots: centre line, median; box, interquartile range; whiskers, min–max. For all plots: n=5 for non-treated, BM-MSCs single and n=6 for BM-MSCs and TAF-MSCs repeated.

**Extended Data Fig. 5:**
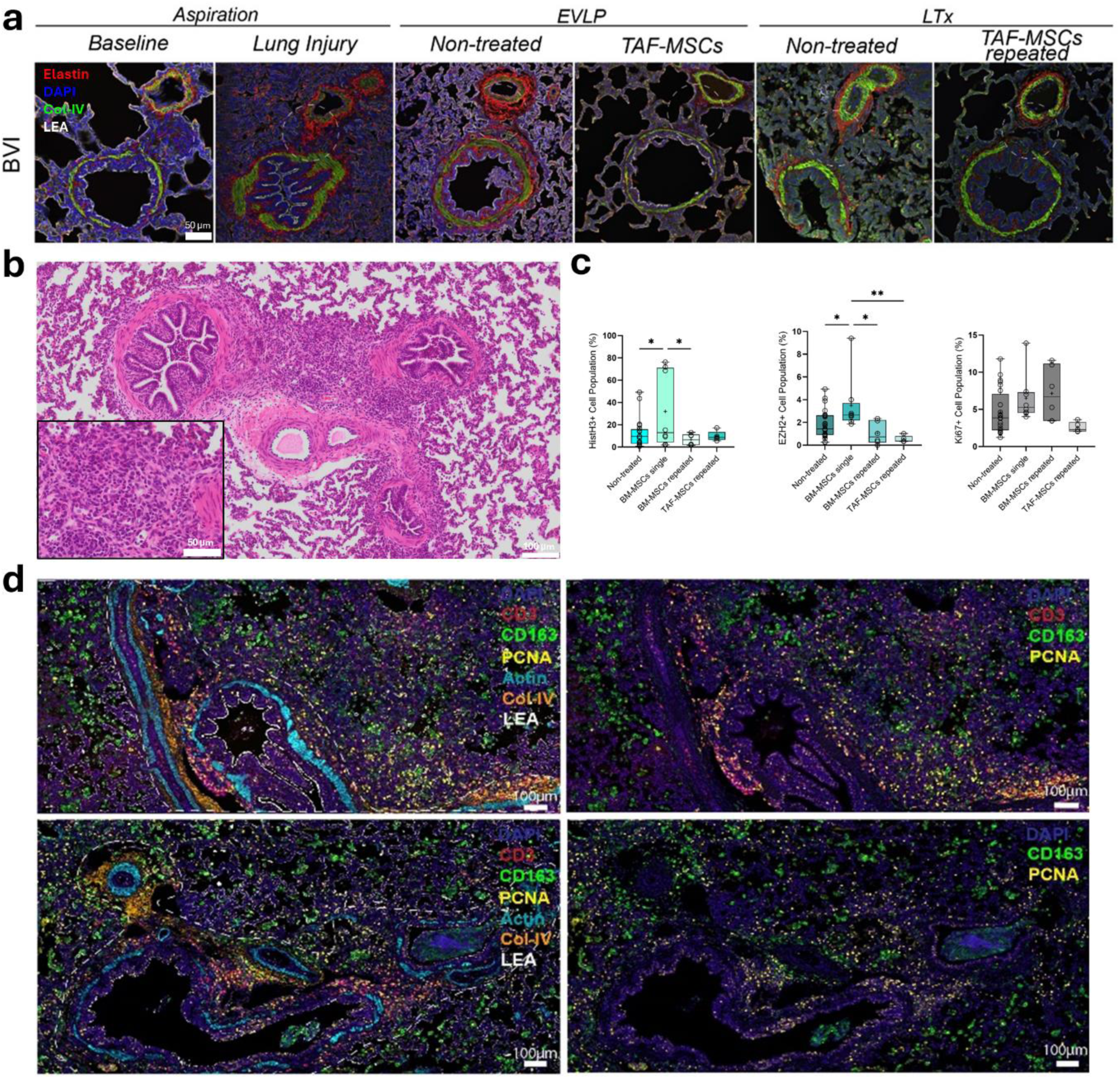
Non-treated grafts displayed substantial immune cell invasion focused at the bronchial-vascular interface (BVI). Throughout the experiment, tissue biopsies were taken and analysed by confocal imaging and standard histological staining. At the endpoint samples were analysed by confocal imaging and 15-plex ultra-high content imaging (MACSima™). **a)** Representative confocal images at baseline, lung injury, EVLP, and post-LTx for repeated dose TAF-MSC and non-treated samples with focus on the BVI. 4′,6-diamidino-2-phenylindole (DAPI), Collagen IV(Col-VI) and Lycopersicon esculentum Lectin (LEA). Scale Bar: 50µm. **b)** Representative image of the BVI in Hematoxylin& Eosin-stained lung tissue in lung injury. Scale bar: 100µm (overview), 50µm (callout). **c)** Quantification of Histone H3+ (HistH3), Enhancer of zeste homolog 2^+^ (EZH2), Antigen Ki67^+^ (Ki67) cell populations (% of whole population) in different regions of interest (ROI) in lung tissue at the endpoint (Non-treated n=23, BM-MSCs single n=8, BM-MSCs repeated n=6, TAF-MSCs repeated n=5). **d)** Representative MACSima™ images of the BVIs from non-treated grafts at the endpoint. 4′,6-diamidino-2-phenylindole (DAPI), Proliferating cell nuclear antigen (PCNA), Collagen IV (Col IV), Lycopersicon esculentum Lectin (LEA). Scale bar: 100µm. Statistical differences were calculated by using ANOVA or Kruskal-Wallis test when data were not normally distributed. *p<0.05, **p<0.01, ***p<0.001, ****p<0.0001. Boxplots: centre line, median; box, interquartile range; whiskers, min-max.

**Extended Data Fig. 6:**
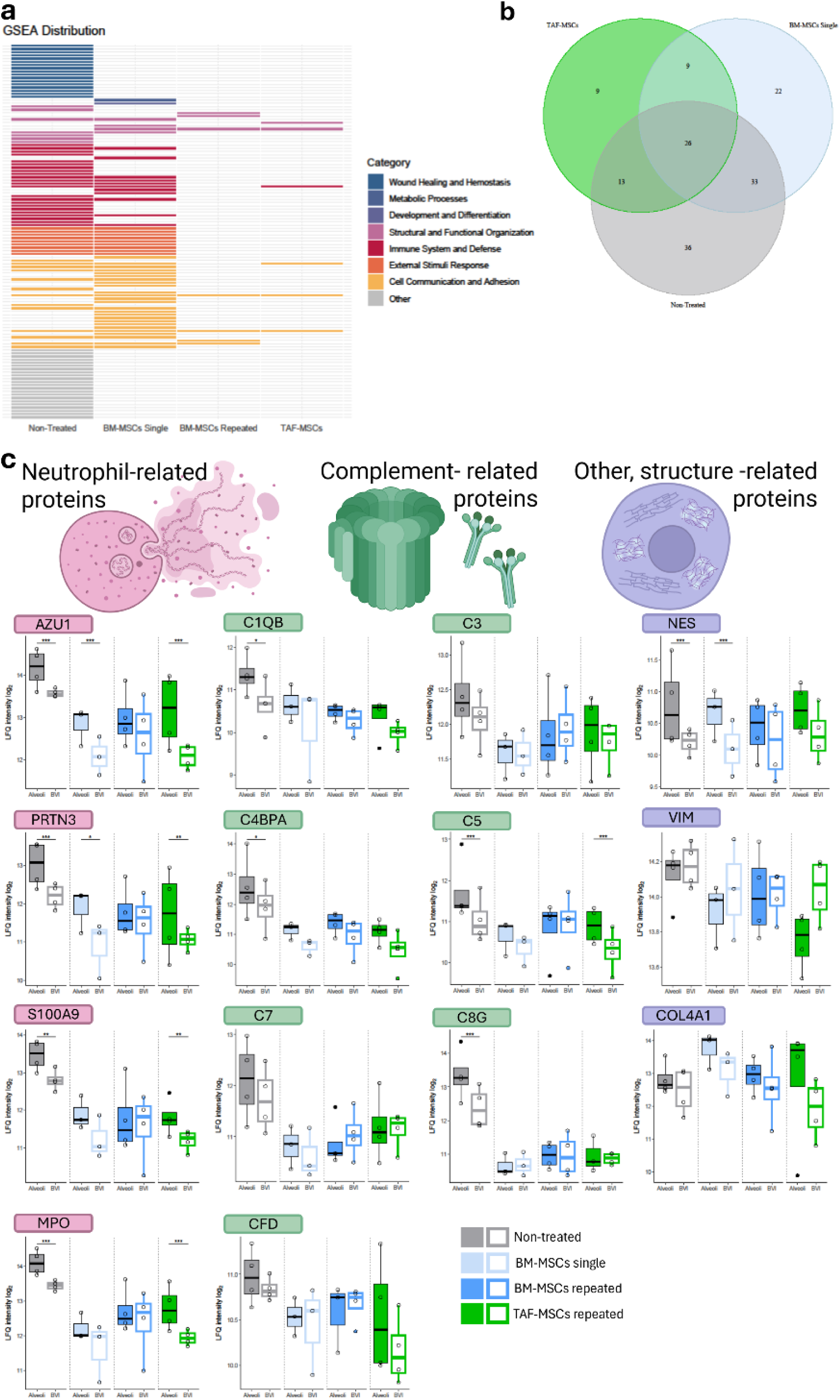
The bronchial-vascular interface (BVI) is an immunological hotspot and differs in proteomic signature to the alveoli. After three days of monitoring and four hours of isolated graft evaluation, tissue from all groups was analysed by mass spectrometry following selective laser capture microdissection of BVI and alveolar regions. n=4 per region and group for non-treated, BM-MSCs and TAF-MSCs repeated and n=3 for BM-MSCs single. **a)** Gene set enrichment analysis (GSEA) displaying differentially expressed pathways in Alveoli versus BVI across groups. **b)** Venn diagram of pathways related to immune system and defence that are differentially expressed between alveoli and BVI. **c)** Quantification of selected protein expression levels associated with neutrophils, the complement cascade, and structural integrity, comparing alveoli and BVI. See Supplementary Table 2 for protein abbreviations. All individual values presented as log_2_ fold change in intensity with FDR corrected p-values (q-values): *p<0.05, **p<0.01, ***p<0.001, ****p<0.0001. Boxplots: centre line, median; box, interquartile range; whiskers, min-max.

