## Supplemental for "Cell therapy for regeneration of injured donor lungs for transplantation"

**Supplementary Methods**

*Animal Preparation*

The pigs utilized in the study had a mean weight of 40 kg. Blood typing was performed prior to the experiment using Seraclone™ Anti-A reagent (blood grouping reagent, Bio-Rad, Medical Diagnostics GmbH, Dreieich, Germany), and the pigs paired into donor and recipients accordingly.

All pigs received premedication including ketamine (Ketaminol® vet. 100 mg/mL; Farmaceutici Gellini S.p.A., Aprilia, Italy; 20 mg/kg) and xylazine (Rompun® vet. 20 mg/mL; Bayer AG, Leverkusen, Germany; 2 mg/kg). Following sedation, a peripheral intravenous (IV) line was placed in the earlobe, and a urinary catheter was surgically inserted into the bladder. General anesthesia was achieved with an infusion of Midazolam (Midazolam Panpharma®, Oslo, Norway), ketamine (Ketaminol® vet, 100 mg/mL; Farmaceutici Gellini S.p.A., Aprilia, Italy; 20 mg/kg) and fentanyl (Leptanal®, Lilly, France) throughout the surgery. Mechanical ventilation was established by using a Servo-i mechanical ventilator (Getinge, Gothenburg, Sweden) following intubation with a 7.5 size endotracheal tube. Ventilation parameters were adjusted accordingly to maintain carbon dioxide levels ($PaCO_2$) between 33-41 mmHg. Tidal volume (Vt) was kept at 6-8 mL/kg. An arterial line (Secalon-T™, Merit Medical Ireland Ltd, Galway, Ireland) was placed into the right common carotid artery and a pulmonary artery catheter (Swan-Ganz CCOmbo V and Introflex, Edwards Lifesciences Services GmbH, Unterschleissheim, Germany) was inserted in the right internal jugular vein.

*Monitoring through hemodynamic and arterial blood gases*

Hemodynamic measurements were performed using an arterial line and thermodilution with a Swan-Ganz catheter. The following haemodynamic parameters were measured and recorded: Oxygen saturation (sat), heart rate (HR), systolic blood pressure (SBP), diastolic blood pressure (DBP), mean arterial pressure (MAP), central venous pressure (CVP), systolic pulmonary pressure (SPP), diastolic pulmonary pressure (DPP), mean pulmonary pressure (MPP), pulmonary artery wedge pressure (PAWP), systemic vascular resistance (SVR), pulmonary vascular resistance (PVR), and cardiac output (CO).

*Ex vivo lung perfusion (EVLP)*

In case of decreased perfusate level, defined as less than 300 mL in the reservoir, additional Steen solution (XVIVO Perfusion) was added. EVLP was performed for four hours, and haemodynamic parameters and arterial blood gases were recorded throughout its duration. Biopsies, bronchoalveolar lavage fluid (BALF) and blood samples were collected (Figure 1). Afterward, the lungs were cooled down to a temperature of 10 °C for approximately 45 minutes prior to the transplantation.

*Treatment with bone-marrow derived and full term-amniotic fluid-derived MSCs*

Human bone marrow (BM) was collected from 20 to 25-year-old healthy consenting volunteers. BM was aspirated as approved by the Swedish Ethical Review Authority at the Haematology Department, Lund, Sweden. The MSCs were purified by density gradient centrifugation (Ficoll-Paque PREMIUM, 1.077 g/ml, GE Healthcare Life Sciences AB, Uppsala, Sweden), further isolated through plastic adherence and expanded in a NUNC cell factory system (Thermo Fisher

Scientific, Waltham, Massachusetts) to facilitate large-scale production. Cells were maintained in culture and propagated using a GMP-compatible animal serum-free culturing protocol using pooled human platelet lysate as described previously[1] and frozen in early passage until use. The cultured MSCs were characterized by their surface marker expression via flow cytometry and differentiation assays, and tested for their immunomodulatory potential[2].

TAF-MSCs were isolated following a previously described protocol[3,4]. Lung-specific TAF-MSCs were isolated based on an RNA and phenotypic expression profile similar to fetal lung MSCs, and after identification of marker genes for this lung-specific type, fluorescence-activated cell sorting was performed to isolate the lung-specific cell product (AmnioPul-02, Amniotics AB, Lund, Sweden). The TAF-MSCs were characterized for a lack of CD45, CD31, and HLA-DR expression and positive expression of CD90, CD73, and CD105. The TAF-MSCs have been used in a Phase 1b (open-label, dose-escalation, safety trial) for the treatment of lower respiratory disease, showing good safety results in the interim analyses without adverse events (Clinical Trial Number: NCT05348772).

*Measurements of cytokine levels in plasma and bronchoalveolar lavage fluid*

Plasma and bronchoalveolar lavage fluid (BALF) for cytokine analysis were collected at the start of the experiment (baseline), after confirmation of lung injury, at the end of EVLP, and at 72 hours after LTx. Cytokine levels of nine selected cytokines (IL-1β, IL-4, IL-6, IL-8, IL-10, IL-12p40, IFN-α, IFN-γ, and TNF-α) were measured using the multiplex kit Cytokine & Chemokine 9-Plex Porcine ProcartaPlex™ Panel 1 (Thermo Fisher Cat. No. EPX090-60829-901, Thermo Fisher Scientific, Waltham, Massachusetts, US). The kit was run according to manufacturer's instructions and analysed using a Bioplex-200 system (BioRad, Hercules, CA, USA).

*Total Bile Acid Assay*

Amount of total bile acids (TBA) was analysed in BALF taken at Baseline and after confirmation of lung injury. Briefly, all samples apart from Baseline samples were diluted 1:4 and assayed according to manufacturer's instructions (Bile Acid Assay Kit (Colorimetric), ab239702, Abcam, Cambridge, United Kingdom). Absorbance was measured at 405 nm for 60 minutes using kinetic measuring mode on a FLUOStar Omega Microplate Reader (BMG Labtech, Ortenberg, Germany) and TBA concentration calculated according to kit instructions.

*Histopathological Analysis*

Biopsies were obtained from the lungs of healthy pigs (baseline), after confirmation of lung injury at the time of lung harvest, at the start and end of EVLP and at termination of the experiment. To avoid sampling bias, all biopsy locations were predetermined prior to the start of the experiment and randomized. All tissue specimens were fixed overnight in 10% formalin solution, neutrally buffered (Sigma Aldrich, Germany), at 4°C. Prior to infiltration with Histowax (Histolab, Askim, Sweden), fixed tissue specimens were dehydrated using a graded ethanol series, followed by xylene and paraffin. Paraffin-embedded specimens were then cut at a thickness of 5 μm on a RM2255 rotary microtome (Leica, Illinois, US) and mounted on glass slides. Dried sections were de-paraffinized and stained with Haematoxylin and Eosin (Histolab) as per standard protocol. All sections were mounted using Pertex (Histolab) and dried overnight at room temperature. Stained specimens were imaged on a VS-120 microscope (Olympus, Tokyo, Japan) using brightfield acquisition in a scanning mode. Acquired images were de-identified, randomized, and sent out for scoring to four independent scorers with extensive experience in the histology of pig lung tissue. The categories scored were: Neutrophils/immune cells in alveolar space; neutrophils/immune cells in interstitial space; proteinaceous debris/fibrin, also in alveolar space; alveolar septal thickening/structural changes and haemorrhage (erythrocytes), hyaline membranes, other damage (enhanced injury). Scoring was performed using a scale of 0-6 with 6 being the most severe (0 = not present; 1 = scattered, light; 2 = medium light; 3 = medium; 4 = medium heavy; 5 = heavy; 6 = severe), with no weighting of the different categories.

*TUNEL Staining*

TUNEL Assay Kit BrdU Red was used (Abcam, ab66110) according to suppliers' instructions. 10µm FFPE lung sections were de-paraffinized and rehydrated according to standard protocols. Rehydrated samples were permeabilised for 10min at room temperature in 3% triton-x-100, after which staining was carried out. Before mounting, samples were counterstained with tomato lectin-488. Imaging was carried out on a Nikon Ti2 epifluorescence microscope. Samples were imaged blinded, and 4 random FOVs were acquired from each biopsy. Analysis was carried out in Fiji, and TUNEL-positive cells were automatically detected using a pre-developed macro.

*SEM Tissue Processing and Imaging*

Tissue was cut into 200 µm sections using a vibrating microtome (Leica VT1200S, Leica, IL, US) as previously described[5]. To provide structural support for sectioning, samples were embedded in 3% low melting point agarose (Sigma-Aldrich, Darmstadt, Germany) in distilled water. Samples were first washed twice in 0.1 M Sorensen´s buffer pH 7.4 to remove media, then fixed in approximately 10 times the sample volume of "SEM fix" (0.1 M Sorensen´s phosphate buffer pH 7.4, 2% formaldehyde, and 2% glutaraldehyde) at room temperature for at least 30 min. After fixation, the samples were washed twice in 0.1 M Sorensen´s buffer, pH 7.4, to remove excess fixative. Samples were then dehydrated in a graded series of ethanol (30%, 50%, 70%, 80%, 90% and twice in 100%) and subsequently critical point dried before being mounted on aluminium stubs, sputtered with 10nm Pt/Pd (80/20) in a Quorum Q150T ES turbo pumped sputter coater. Imaging was done on a Jeol JSM-7800F FEG SEM (JOEL, Tokyo, Japan).

*Confocal Tissue Processing and Imaging*

Ten µm FFPE lung sections were de-paraffinized and rehydrated according to standard protocols. Antigen retrieval was carried out in citrate buffer for 30 min at 98°C. Samples were permeabilized in 0.5% triton-x-100, 1% normal goat serum, and 5% bovine serum albumin. Primary antibodies were incubated overnight at 4°C in PBS (PCK-1:500, Elastin-1:200, SMA-1:500). Secondary antibodies were incubated for 90 min at 4°C (goat anti-rabbit 568, goat anti-mouse-647). Finally, samples were incubated for 30 min with DAPI (1:1000) and tomato lectin-488 (1:500). Mounting took place with fluoromount-G and 1.5 mm glass coverslips. Image acquisition took place on a Nikon A1RHD equipped with a 20x objective (NA-0.8) and piezo 200 stage.

*Ultra-High-content Tissue Processing and Imaging*

Formalin-fixed paraffin-embedded (FFPE) lung sections (10 µm) were deparaffinized, rehydrated, and subjected to antigen retrieval in TEC buffer at 98°C for 30 min. Sections were prestained with DAPI (1:10, 10 min) and washed in running buffer before imaging. Each well was filled with 250 µL running buffer prior to loading into the MACSima™ Imaging Platform with MACSwell 4 frames (Miltenyi Biotec, Bergisch Gladbach, Germany), as previously described[6]. Antibodies included both pre-validated reagents from Miltenyi Biotec and externally sourced antibodies, all directly conjugated to FITC, PE, APC, Alexa Fluor 488, or Alexa Fluor 635. Staining parameters (dilution, incubation, cycle sequence, photobleaching, and exposure) were standardized across experiments. Image acquisition was performed with a 20× long working distance objective (NA 0.45).

*Proteomic analysis with mass spectrometry*

Protein was extracted from homogenized tissue and solubilized in 2% SDS and subsequently digested with an S-TRAP protocol. Reduction was performed with 20 mM dithiothreitol (DTT) for 45 minutes at 56°C and then 40 mM iodoacetamide (IAA) in the dark at room temperature for 30 minutes. Samples were acidified with 2.5% phosphoric acid and washed with buffer before binding to an S-Trap micro column (ProTifi, New York, United States). Samples were double digested overnight at 37°C with lysine-C (at a 1:50 ratio of enzyme to protein by ng) and trypsin (at a 1:50 ratio of enzyme to protein by ng). Peptide fractionation, data-dependent acquisition (DDA), and data-independent acquisition (DIA) was performed as previously reported[4] using a TIMS TOF HT instrument.

Raw LC-MS/MS data were analysed using DIA-NN v 1.8.1 with trypsin set as enzyme and with the following parameters: high_precision: true, global_normalisation: false, library_free: true, missed cleavages: 2. The FASTA file was downloaded from the Uniprot database with identifier ID: UP000008227_9823 and FDR set at 1%. Subsequently, the output files were loaded into RStudio v 2022.12.0 with R v 4.3.1. The MS-DAP package[7] was used for normalization and differential expression analysis using the following parameters: minimum detected peptides: 1, filtered at 75% identified proteins per contrast. Quality control analysis revealed two samples, one from the BM-MSC single dose and one from the BM-MSC repeated dose group, with low peptide identifications, and were thus removed from further downstream analysis.

Normalization was performed using variance stabilizing normalization. Differential expression was performed using the MS-EmpiRe R package[8]. Significantly differentially expressed proteins were defined as FDR corrected p-values (q-values) > 0.05 and Log 2-fold cutoff values defined by bootstrapping. For the heatmap, MaxLFQ values were normalized using Z-scores and plotted using the heatmap package v 1.0.12 with Euclidean clustering. GO term enrichment analysis was performed using geneontology.org's PANTHER GO Enrichment Analysis using the overrepresentation test on all identified proteins in at least 75% of samples.

For the microdissection, the same tissue blocks used for the confocal and MACSima image processing were utilized, and tissue sections were placed on PEN membrane slides. These slides were then deparaffinized, and regions of interest, including alveolar regions and the bronchial-vascular interface (BVI), were identified by direct visualization. Laser-capture microdissection (LCM) was performed on a PALM MicroBeam system (Carl Zeiss MicroImaging, Jena, Germany) with a 10x objective using the PALM Robo 4.5 Pro Software (Carl Zeiss MicroImaging) with the Robo-LAP mode, by which defined regions were cut out and lifted in one piece by the laser. Dissected material was catapulted onto the lids of adhesive cap tubes (Zeiss, Jena, Germany), and the CapCheck function enabled inspection of the collected tissue on the cap lid. An average area of 1.0 x $10^6$ $mm^2$ was collected from a section of alveolar tissue and from a section of BVI per sample. The collected tissue was then processed and analysed by mass spectrometry using the same methods as outlined above. Quality control analysis revealed two samples from the same animal in the BM-MSC single-dose group with low peptide identifications, which were removed from further downstream analysis.

### Supplementary Tables

**Supplementary Table 1: Clinically relevant parameters during 0-60 hours follow-up.** Clinically relevant measurements in the post-transplantation follow-up including the non-treated (bold, n=6), the BM-MSC single dose (n=6, light grey), the BM-MSC repeated dose (n=6, medium grey) or the TAF-MSC repeated group (n=6, dark grey). All parameters are given as mean ± standard deviation: oxygen saturation (Sat, %), heart rate (HR, beats per minute, bpm), systolic blood pressure (SBP, mmHg), diastolic blood pressure (DBP, mmHg), mean arterial pressure (MAP, mmHg), central venous pressure (CVP, mmHg); systolic pulmonary pressure (SPAP, mmHg), diastolic pulmonary pressure (DPAP, mmHg), mean pulmonary pressure (MPP, mmHg), pulmonary artery wedge pressure (PAPWP, mmHg), cardiac output (CO, L/min), cardiac index (CI, L/min/$m^2$), systemic vascular resistance (SVR, dynes*s/$cm^5$), pulmonary vascular resistance (PVR, dynes*s/$cm^5$), systemic vascular resistance index (SVRI, dynes*s/$cm^5$*$m^2$), pulmonary vascular resistance index (PVRI, dynes*s/$cm^5$*$m^2$), mechanical ventilator settings with volume-controlled ventilation: minute volume (MV, L/min), peak inspiratory pressure (PIP, $cmH_2O$), peak end expiratory pressure (PEEP, $cmH_2O$), tidal volume (Vt, mL), respiratory rate (RR, breaths/min), pH, partial pressure of carbon dioxide ($PaCO_2$, mmHg), haemoglobin (Hb, g/L), lactate (mmol/L), base excess (BE, mmol/L).

| | Baseline | 1h | 12h | 24h | 36h | 48h | 60h |
|---|---|---|---|---|---|---|---|
| **Sat (%)** | **97.5±1.4** | **97.5±0.5** | **97.7±1.2** | **98±0.9** | **97.7±1.4** | **98±1.7** | **98±1.2** |
| | 98.7±1 | 96.8±1.2 | 96.5±1.8 | 95.3±2.4 | 98.3±1 | 97±1.3 | 96.5±3.3 |
| | 99.3±0.5 | 97.8±1.2 | 98±1.3 | 97.8±1.2 | 97.8±1 | 97.8±0.4 | 98.2±1.2 |
| | 99.8±0.4 | 98.3±1.2 | 98.5±0.5 | 97.5±1.8 | 97.8±1.7 | 97.5±0.8 | 97±1.1 |
| **HR (bpm)** | **82±17.7** | **74.5±7.5** | **78.8±19.5** | **76.2±10.6** | **78.5±10.8** | **77±18.1** | **72.4±13.9** |
| | 84.3±25.5 | 82.5±16.3 | 81.8±11.7 | 90.3±23.6 | 82.7±22.9 | 91.7±24.6 | 87.5±27.8 |
| | 85.7±18.1 | 77.8±9.4 | 76.2±11.8 | 84.5±17 | 78.3±4.8 | 75.5±9 | 76.7±9.8 |
| | 87.5±10.6 | 87.7±11.7 | 73.3±13.3 | 83.5±11.3 | 79.3±10.1 | 81±13.4 | 89.5±22 |
| **SBP (mmHg)** | **117.3±12.6** | **112.3±9** | **108.2±11.5** | **107±6.8** | **109.8±4.6** | **108.3±8.8** | **112.8±13.3** |
| | 116.8±6.4 | 113.8±12.7 | 104.7±8.1 | 104.5±6.9 | 108±9.7 | 113.2±5.8 | 112.5±11.2 |
| | 110.2±23.8 | 114.5±9.1 | 112.8±8.4 | 106.7±6.2 | 113.8±7 | 116.5±14.4 | 118.2±10.7 |
| | 109.3±9.6 | 117.8±12.9 | 105±5.6 | 108.3±8.3 | 113.2±11.5 | 118.3±14.1 | 110.3±9.9 |
| **DBP (mmHg)** | **74.8±19.1** | **76.7±6.9** | **70.3±10.5** | **71±13** | **78.2±14.6** | **78.5±11.6** | **76.4±13.5** |
| | 79.8±6.7 | 72±12.1 | 63.5±10.3 | 60.5±7.4 | 62.8±13.4 | 67.5±12.7 | 67.5±10.3 |
| | 73±14.9 | 76.7±5.5 | 74.3±7.3 | 68±5.7 | 80±10.5 | 81.3±12.7 | 80±12.5 |
| | 72±5.6 | 77.2±11.5 | 67.3±9.3 | 59±11 | 67.5±14.2 | 75.2±11.6 | 69.3±8.3 |
| **MAP (mmHg)** | **88.8±17.5** | **94.3±5.2** | **89.2±9.5** | **89±10.5** | **95.7±11** | **95.8±9.7** | **94±12** |
| | 96.3±7.2 | 89.8±13.2 | 80.7±10.4 | 77.5±8.1 | 80.3±13.6 | 86.5±10.6 | 85.8±10.3 |
| | 88.5±18.2 | 94.7±8.4 | 92.7±7.3 | 86.2±7.3 | 96.2±7.5 | 100.5±17.8 | 99.7±11.7 |
| | 87.7±7.2 | 93.8±12 | 85±8.7 | 79.5±10.6 | 87±14.1 | 93.5±12.4 | 86.5±9.2 |
| **CVP (mmHg)** | **7.2±1.5** | **4.8±2.8** | **6.7±2.5** | **5±3.1** | **7.2±2.8** | **6.2±2.9** | **7±1.8** |
| | 7.3±2.4 | 6.5±1.9 | 5.5±2 | 7±2.5 | 7.5±2.9 | 8.3±2.6 | 7.2±2.9 |
| | 6.7±3.3 | 7.7±2.5 | 6.7±2.3 | 5.8±1.2 | 7.2±1.5 | 8±4.5 | 5.7±0.8 |
| | 5±2.2 | 5.8±3.3 | 4.7±0.8 | 7.5±4.9 | 6.2±3.3 | 7.7±4.4 | 6.8±1.7 |
| **SPAP (mmHg)** | **23.8±3.4** | **30.3±3.8** | **26.2±7** | **25.3±8.8** | **27.2±7.4** | **29.8±6** | **27.6±6.7** |
| | 26.8±4.4 | 37±4.6 | 31±1.5 | 34.3±5.6 | 34±5.9 | 34.7±6.3 | 35.8±7 |
| | 21.5±2.1 | 33.8±3.4 | 26.3±5.6 | 29.3±4.6 | 29.6±5 | 28±6.4 | 28.3±6.9 |
| | 24.2±6.3 | 35.3±5.1 | 27.7±3.2 | 31.2±2.3 | 31.4±5.3 | 32.8±3.8 | 30.2±4.8 |
| **DPAP (mmHg)** | **14.5±3.9** | **16.2±3.1** | **14±4.4** | **14.2±2.9** | **15.5±3.8** | **12.8±4.7** | **13.4±5.2** |
| | 14.8±2.5 | 19.2±1.7 | 16.2±4.9 | 18.2±6.3 | 17±5.8 | 20±6 | 19.8±6 |
| | 12.7±2.9 | 18.3±3.4 | 13.2±4.6 | 15.2±3.4 | 12.4±3.4 | 14±4 | 15.8±3.1 |
| | 14.3±5.1 | 19.8±4.8 | 13±2.8 | 12.8±2.3 | 13±1.9 | 14.8±3.3 | 14±3 |
| **MPP (mmHg)** | **18.3±3.1** | **22.3±3.1** | **16.2±4.2** | **18±5** | **20±4.8** | **19.5±5.4** | **19±4.6** |
| | 20.5±3.4 | 26.5±1.6 | 22.5±3.3 | 23.8±5.9 | 23.8±5.9 | 25.5±5.4 | 26±6 |
| | 16.3±2.4 | 25±4 | 19.2±4.5 | 21±3.5 | 20±3.2 | 19.8±3.6 | 21±4.8 |
| | 19.3±5.2 | 26±4.7 | 19.3±2.4 | 21±2.1 | 21±2.9 | 21.4±3.2 | 20.8±4.1 |
| **Wedge (mmHg)** | **13.2±2.7** | **11±5.3** | **8.8±1.7** | **8.8±1.3** | **11.2±2** | **11±3** | **9.8±2.5** |
| | 10.2±2.3 | 9.8±1.7 | 10.3±7.1 | 10.5±3.1 | 10.3±3 | 11.8±2.6 | 11.2±3.7 |

| | | | | | | | |
|---|---|---|---|---|---|---|---|
| | 8.8±3.4 | 9±1.5 | 8.3±4.3 | 9.8±1.9 | 10.2±0.8 | 11±3.6 | 10.3±2.7 |
| | 9.2±2.9 | 9.5±4.4 | 9.2±2.1 | 9.2±1.7 | 9.2±2.3 | 10±3.6 | 9.8±2.8 |
| **CO (L/min)** | **4.2±1.3** | **3.6±0.6** | **4.3±0.8** | **5±1.1** | **4.5±0.9** | **4.7±0.8** | **4±0.5** |
| | 4.5±1.1 | 4.4±1.2 | 4.4±0.3 | 5.1±1.2 | 4.4±0.8 | 5±1.5 | 5±1.5 |
| | 3.8±0.8 | 4±1 | 4.8±0.7 | 5.2±1.2 | 4.3±0.5 | 4.1±0.8 | 4.4±0.5 |
| | 5±1.1 | 4.6±0.7 | 4.2±1.1 | 6.2±0.9 | 5.8±0.6 | 5.3±0.3 | 5.6±0.6 |
| **CI (L/min/m²)** | **3.4±1.2** | **2.8±0.3** | **3.5±0.6** | **4±0.9** | **3.6±0.6** | **3.7±0.8** | **3.2±0.4** |
| | 3.6±0.9 | 3.5±0.9 | 3.5±0.3 | 4.2±1 | 3.5±0.7 | 4±1.2 | 4±1.3 |
| | 3±0.6 | 3.2±0.8 | 3.9±0.5 | 4.2±1 | 3.5±0.4 | 3.2±0.4 | 3.6±0.3 |
| | 4±0.9 | 3.7±0.5 | 3.4±0.9 | 4.9±0.7 | 4.6±0.4 | 4.2±0.2 | 4.5±0.4 |
| **SVR (DS/cm⁵)** | **1887.5±446** | **1937.3±338.9** | **1510±533.7** | **1305.8±210.4** | **1704.2±484.9** | **1594.7±354.6** | **1755±313.1** |
| | 1609.7±310.6 | 1649.8±431.2 | 1402.5±268.6 | 1193±362.7 | 1491.8±0 | 1428.2±577.4 | 1473.3±596.5 |
| | 1880.8±498.6 | 1844.3±330.9 | 1480.2±188.3 | 1398.3±248.9 | 1950±0 | 1450±59.4 | 1766±219 |
| | 1326.5±238.7 | 1564±292.7 | 1520.8±494 | 937.3±204.6 | 1160.8±0 | 1375.4±254.5 | 1141±148.3 |
| **PVR (DS/cm⁵)** | **153.8±60.7** | **304.7±87.9** | **188.6±62.7** | **151±66.9** | **176.4±76** | **172.2±67.7** | **190.4±45.3** |
| | 165.8±56.9 | 330.7±91.1 | 219.2±86.4 | 217.3±93.3 | 260±0 | 208±58.5 | 221.2±31.8 |
| | 162.7±43.4 | 347.2±111.2 | 191.8±57.1 | 177.3±24.1 | 219±0 | 183.5±67.2 | 198.8±74.3 |
| | 169.5±58 | 305.7±71.9 | 196.5±63.1 | 158.7±38.3 | 160.8±0 | 170.2±34.7 | 146.8±25.4 |
| **SVRI (DS/cm⁵/m²)** | **2434.7±761.9** | **2468.7±367.9** | **1855±625.2** | **1815.5±415.7** | **2067.4±454.6** | **2034.8±547.1** | **2297±556.8** |
| | 2021.2±416.6 | 2070.8±557.7 | 1760.5±355.7 | 1480.5±482.1 | 1883.6±0 | 1797.8±749.4 | 1854.3±769.7 |
| | 2321.7±558 | 2296.3±454.8 | 1839.3±248.6 | 1662.3±281.9 | 2262±0 | 2203±14.1 | 2167.7±301.3 |
| | 1662.3±300.8 | 1965.3±380.3 | 1936.5±673.8 | 1177.5±256.9 | 1452±0 | 1722±322.2 | 1426.4±166.5 |
| **PVRI (DS/cm⁵/m²)** | **200.8±101.5** | **434.5±280.1** | **232.2±72.6** | **191.3±80.7** | **219±83.7** | **217.5±82.8** | **250±75.8** |
| | 208.2±72.6 | 414.3±114.4 | 274.8±109.1 | 269.7±118 | 326.8±0 | 260.8±74.9 | 277.3±38.2 |
| | 200.7±47.5 | 433.8±152.9 | 239.5±72.8 | 224.2±38.2 | 254±0 | 248.5±128 | 243.8±91.5 |
| | 211.8±70.2 | 384±89.7 | 248±81.1 | 199.7±49.4 | 201.2±0 | 213±42.7 | 182.8±30.6 |
| **MV (L/min)** | **6.2±0.7** | **7.4±0.6** | **8±1.1** | **7.8±1.3** | **7.7±1.3** | **8.1±1.4** | **8.2±1.4** |
| | 6.1±0.6 | 7.5±0.6 | 7.7±0.6 | 7.6±0.8 | 7.8±1 | 7.7±0.8 | 7.9±1.2 |
| | 5.7±0.6 | 7.2±0.9 | 7.5±0.8 | 7.6±0.5 | 7.6±0.6 | 7.7±0.8 | 7.8±0.7 |
| | 6.1±0.8 | 7.1±0.6 | 7.4±0.8 | 7.5±0.5 | 7.5±0.5 | 7.4±0.7 | 7.5±0.6 |
| **PIP ($cmH_2O$)** | **17.8±2.6** | **25.6±1.7** | **24.1±2.1** | **22.7±2** | **23.3±1.7** | **25.1±1.3** | **25.5±3** |
| | 19.7±1.6 | 26.4±2.1 | 24.6±3.3 | 25.6±4.5 | 25.5±4.4 | 25.6±4.9 | 26.9±7.5 |
| | 17.3±1.4 | 24.9±1.7 | 24.4±1.8 | 25.2±2.2 | 25.5±2.7 | 27.5±4.3 | 27.1±3.2 |
| | 17.2±1.9 | 23.3±1.2 | 20.8±1.2 | 21.8±2.7 | 21.5±2.3 | 22.5±3.2 | 22.5±3.1 |
| **PEEP ($cmH_2O$)** | **5.3±0.5** | **5.7±1.2** | **6.2±1.2** | **6.2±1.2** | **5.7±1.2** | **6.7±1.5** | **6.8±1.6** |
| | 5±0 | 5±0 | 5±0 | 6±1.7 | 6.3±1.8 | 5.7±1.4 | 5.2±0.8 |
| | 5±0 | 6±1.5 | 5.5±1.4 | 5.8±1.3 | 6±1.1 | 5.7±1.2 | 5.7±0.8 |
| | 5±0 | 5±0 | 4.8±0.4 | 5.5±1.2 | 5.3±1 | 5.5±1 | 5.7±0.8 |
| **Vt (mL)** | **299.3±38.1** | **282.8±26.6** | **297.8±23.1** | **279.2±10.7** | **286.8±17.6** | **288±15.2** | **303.4±29.1** |
| | 298.8±21 | 279±12.8 | 274.5±18.2 | 281.5±21.3 | 287.8±20.5 | 285.2±20.8 | 278.7±39.4 |
| | 283.7±14.8 | 269.7±16.3 | 270.3±15.4 | 269.2±17.4 | 271.5±11.5 | 271±17.4 | 266.8±22.7 |
| | 311.7±38.2 | 298.3±24.8 | 307.8±21.3 | 311.2±25.1 | 309.5±24.7 | 318.5±21.6 | 322.5±26.4 |
| **RR (breaths/min)** | **21±1.7** | **26.2±2.1** | **27±3.2** | **27.3±3.1** | **27.3±3.1** | **27.8±3.3** | **26.9±2.7** |
| | 20±1.1 | 26.5±3.6 | 28±2.6 | 27.8±2.8 | 27.9±3 | 27.7±2.2 | 28±2.4 |
| | 20.5±2.8 | 26.8±1.9 | 28±2.5 | 28.4±2.1 | 28.3±2.2 | 28.5±1.6 | 29.5±1.6 |
| | 19.3±2.7 | 22.8±2.3 | 23.5±2.1 | 23.5±2.2 | 25.3±4 | 24.8±2.7 | 25±2.7 |
| **pH** | **7.5±0.1** | **7.4±0** | **7.4±0.1** | **7.4±0** | **7.5±0** | **7.4±0** | **7.5±0** |
| | 7.5±0 | 7.4±0 | 7.4±0 | 7.4±0.1 | 7.5±0 | 7.5±0 | 7.4±0.1 |
| | 7.5±0 | 7.4±0 | 7.4±0 | 7.4±0 | 7.4±0 | 7.5±0 | 7.4±0 |
| | 7.4±0.1 | 7.4±0 | 7.4±0 | 7.4±0 | 7.5±0 | 7.5±0 | 7.5±0 |
| **$PaCO_2$ (mmHg)** | **5.9±0.8** | **6.4±0.7** | **6±0.7** | **6.2±0.5** | **6.2±0.4** | **6.5±0.7** | **6±0.7** |
| | 6±0.5 | 6.7±0.3 | 6.1±0.5 | 6.4±1 | 6.3±0.5 | 6.5±0.5 | 6.7±1.1 |
| | 5.9±0.7 | 6.7±1 | 6.4±0.6 | 6.5±0.4 | 6.4±0.4 | 6.2±0.4 | 6.3±0.5 |
| | 5.9±0.9 | 6.4±0.5 | 5.9±0.3 | 6.2±0.2 | 6.2±0.5 | 5.8±0.5 | 6±0.4 |
| **Hb (g/L)** | **106.7±12.9** | **108.3±10.7** | **101.5±14** | **90.3±12.4** | **86.7±11.8** | **84.8±11.5** | **86.2±8.5** |
| | 108.7±4.9 | 104.7±19.2 | 89.8±17.1 | 76±11.8 | 71.7±15.6 | 67.7±20.3 | 66.3±19 |
| | 111.5±9.2 | 108.5±12.6 | 94.3±8.1 | 86.5±6.9 | 86.7±11.3 | 83.5±10.3 | 82±9.3 |
| | 111.8±9.1 | 115.3±7.3 | 101.3±7.4 | 88±8.6 | 88±8.5 | 84.7±4.6 | 81.3±4.7 |
| **Lactate (mmol/L)** | **1.2±0.3** | **1.1±0.4** | **1.1±0.9** | **0.8±0.6** | **0.5±0.1** | **0.6±0.1** | **0.6±0.1** |
| | 1.1±0.3 | 1.3±0.4 | 0.8±0.4 | 0.8±0.3 | 0.7±0.2 | 0.7±0.2 | 0.9±0.3 |
| | 1±0.2 | 2±2.1 | 0.8±0.6 | 0.6±0.2 | 0.5±0.1 | 0.6±0.2 | 0.6±0.2 |

| | | | | | | | |
|---|---|---|---|---|---|---|---|
| | 1.6±1.2 | 1.2±0.2 | 1.1±0.4 | 0.7±0.2 | 0.7±0.4 | 0.7±0.2 | 0.8±0.2 |
| **BE (mmol/L)** | **6.7±1.9** | **5.2±2.4** | **5.5±4.7** | **8.3±3.3** | **10.2±5.1** | **9.7±6.3** | **9.2±5.4** |
| | 7.7±1.1 | 8.1±2.3 | 7±2.8 | 7.4±2.3 | 10.3±1.8 | 10.5±3 | 10.1±5.2 |
| | 7.4±1.1 | 3.3±4 | 6.4±3.8 | 7.1±2.6 | 9.1±2.6 | 9.8±3.5 | 7.7±2.1 |
| | 5.6±2.8 | 5.5±2.9 | 6.2±3.8 | 6.2±3.1 | 10±3.4 | 9.8±2.4 | 7.9±3.5 |

**Supplementary Table 2: Abbreviations for relevant proteins from mass spectrometry analysis.** After three days monitoring and four hours isolated graft evaluation, tissue from all groups was analysed by mass spectrometry, either directly on whole tissue pieces or following selective laser capture microdissection of BVI and alveolar regions. This table presents a summary of all relevant proteins mentioned throughout the manuscript.

| Gene name | Protein name | UniProt ID | Whole tissue proteomics | | Microdissection proteomics | |
|---|---|---|---|---|---|---|
| | | | Detected | Graph placement | Detected | Graph placement |
| **AIF1** | Allograft inflammatory factor 1 | P81076 | Yes | Extended Fig. 4 | No | / |
| **AZU1** | Azurocidin | P80015 | Yes | Extended Fig. 4 | Yes | Extended Fig. 6 |
| **C1QB** | Complement C1q B chain | A0A5G2QNN2 | Yes | Extended Fig. 4 | Yes | Extended Fig. 6 |
| **C1QC** | Complement C1q C chain | A0A286ZSJ7 | Yes | Fig. 5 | No | / |
| **C4BPA** | Sushi domain-containing protein | F1S0J2 | Yes | Extended Fig. 4 | Yes | Extended Fig. 6 |
| **C5** | Complement C5 | A0A287AIM8 | Yes | Extended Fig. 4 | Yes | Extended Fig. 6 |
| **C5AR1** | C5a anaphylatoxin chemotactic receptor 1 | I3LUE7 | Yes | Extended Fig. 4 | No | / |
| **C6** | Complement component C6 | A0A5G2R8T9 | Yes | Extended Fig. 4 | No | / |
| **C7** | Complement component C7 | Q9TUQ3 | Yes | Fig. 5 | Yes | Extended Fig. 6 |
| **C8G** | Complement C8 gamma chain | A0A287AFQ4 | Yes | Extended Fig. 4 | Yes | Extended Fig. 6 |
| **C9** | Complement C9 | A0A8W4F8A6 | Yes | Extended Fig. 4 | Yes | Fig. 6 |
| **CD14** | Monocyte differentiation antigen CD14 | A2SW51 | Yes | Fig. 5 | Yes | Fig. 6 |
| **CD163** | Scavenger receptor cysteine-rich type 1 protein M130 | Q2VL90 | Yes | Fig. 5 | Yes | Fig. 6 |
| **CD3E** | T-cell surface glycoprotein CD3 epsilon chain | Q7YRN2 | Yes | Fig. 5 | No | / |
| **CD4** | T-cell surface glycoprotein CD4 | I3L556 | Yes | Fig. 5 | No | / |
| **CD40** | Tumor necrosis factor receptor superfamily member 5 | Q8SQ34 | Yes | Extended Fig. 4 | No | / |
| **CFD** | Complement factor D | P51779 | Yes | Fig. 5 | Yes | Extended Fig. 6 |
| **CFI** | Complement factor I | A0A287AQ20 | Yes | Extended Fig. 4 | No | / |
| **COL4A1** | Collagen type IV alpha 1 chain | A0A8W4FCC2 | Yes | Extended Fig. 4 | Yes | Extended Fig. 6 |
| **MPO** | Myeloperoxidase | K7GRV6 | Yes | Fig. 5 | Yes | Extended Fig. 6 |
| **NES** | Nestin | I3LNY6 | Yes | Extended Fig. 4 | Yes | Extended Fig. 6 |
| **OLR1** | Oxidized low-density lipoprotein receptor 1 | Q9TTK7 | Yes | Fig. 5 | No | / |
| **PRTN3** | Proteinase 3 | A0A8D0VTQ5 | Yes | Fig. 5 | Yes | Extended Fig. 6 |

| | | | | | | |
|---|---|---|---|---|---|---|
| **S100A8** | Protein S100 | C3S7K5 | Yes | Fig. 5 | Yes | Fig. 6 |
| **S100A9** | S100 calcium binding protein A9 | K7GME6 | Yes | Extended Fig. 4 | Yes | Extended Fig. 6 |
| **SELL** | L-Selectin | A0A288CG53 | Yes | Fig. 5 | No | / |
| **VIM** | Vimentin | P02543 | Yes | Extended Fig. 4 | Yes | Extended Fig. 5 |
